# MRSIPrep: A Standardized Post-Quantification Framework for Preprocessing Whole-Brain Magnetic Resonance Spectroscopic Imaging

**DOI:** 10.64898/2026.09.11.750896

**Authors:** Federico Lucchetti, Edgar Céléreau, Raoul Jenni, Jean-Baptiste Ledoux, Stephan Eliez, Farnaz Delavari, Patric Hagmann, Yasser Alemán-Gómez, Antoine Klauser, Paul Klauser

## Abstract

Magnetic resonance spectroscopic imaging (MRSI) enables non-invasive mapping of neurometabolites levels across the human brain. Although spectral fitting and metabolite quantification are increasingly supported by mature software tools, the downstream processing of quantified metabolite maps remains heterogeneous across laboratories. Here, we introduce MRSIPrep, an open-source, modular, and reproducible post-quantification framework for whole-brain MRSI. MRSIPrep standardizes quality control, tissue correction, spatial normalization, atlas projection, and derivative generation from quantified metabolite maps and associated quality metrics. The framework produces voxelwise, regional, and connectomics-ready outputs together with automated quality-control reports. We describe the architecture of MRSIPrep and demonstrate its utility for reproducible MRSI analysis across datasets, acquisition protocols, and downstream applications.

## 1 Introduction

Magnetic resonance spectroscopic imaging (MRSI) enables the non-invasive mapping of multiple neurometabolites across large portions of the human brain, providing a unique window into the metabolic organization of neural systems. Whole-brain proton MRSI was established as feasible for population-level metabolite mapping nearly two decades ago [1], and the last ten years have seen a wave of acceleration strategies to substantially improve its spatial coverage, resolution, and quantification reliability [2, 3]: shortened repetition times as in free-induction-decay (FID) MRSI, Cartesian and non-Cartesian spatial-spectral encoding (e.g. EPSI [2], spiral [4], concentric-ring trajectories as in ECCENTRIC [5]), k-space/time-domain undersampling via parallel imaging and compressed sensing, and subspace/low-rank reconstruction from spatial-spectral priors (e.g. SPICE [6, 7]), comprehensively reviewed by Bogner et al. [3]. Modern techniques increasingly combine several of these strategies at once, as in ECCENTRIC’s pairing of circular trajectories, compressed sensing, FID acquisition, and low-rank reconstruction [5]; as this combined acceleration keeps improving acquisition time, resolution, and SNR, we expect these once-specialized techniques to see substantially wider adoption. These developments have expanded the application of MRSI from regional investigations to studies of large-scale brain organization [8], network architecture, and disease-related alterations [9, 10].

Despite these advances, downstream processing of whole-brain MRSI remains fragmented across laboratories. Mature tools exist for spectral reconstruction and quantification, including LCModel [11], TARQUIN [12], Osprey [13] and FSL-MRS [14], but no widely adopted framework standardizes the subsequent processing of quantified metabolite maps. MIDAS [15] provides an important precedent for whole-brain MRSI, spanning reconstruction, spectral processing, quality control, and metabolite-map analysis, but has principally been developed around EPSI workflows. MRS4Brain [16] provides a related integrated framework for preclinical MRSI. Outside such systems, laboratories commonly rely on custom workflows for quality control, tissue correction, spatial normalization, atlas projection, and feature extraction, limiting reproducibility, cross-site comparability, hindering data sharing and scientific collaboration.

The MRS/MRSI community has already recognized this standardization gap before, though mostly for the acquisition side of the problem: the Minimum Reporting Standards for in vivo Magnetic Resonance Spectroscopy (MRSinMRS) consensus [17] established a community-wide checklist of sequence, hardware, and reconstruction parameters that studies should report, precisely because undocumented acquisition and post-processing choices had been found to materially affect interpretability and reproducibility across the field. MRSinMRS addresses what authors must report about an acquisition; it does not, by itself, constrain or standardize what happens to the data afterward. The processing side of that same problem is what MRSIPrep aims to close, and, as described below (§2.3.3), MRSIPrep integrates MRSinMRS directly into its own quality-control reports so that a study’s acquisition parameters and its processing provenance are documented side by side rather than in separate, disconnected artifacts.

The growing interest in network-based and multimodal analyzes further amplifies these challenges especially if modern neuroimaging hopes to integrate MRSI with structural, functional, molecular, and clinical data, which requires standardized representations that can be compared across cohorts and institutions. In particular, emerging applications such as metabolic connectomics, gradient analysis [8], and machine-learning-based patient stratification depend critically on consistent preprocessing procedures and transparent quality assessment. However, unlike other neuroimaging modalities, for which robust community standards have emerged, the MRSI community lacks a unified framework that bridges the gap between metabolite quantification and downstream neuroscientific analysis, of the kind that fMRIPrep and the broader NiPreps community have established for structural or functional MRI. [18, 19].

To address this need, we introduce MRSIPrep, an open-source, reproducible, and modular framework for post-quantification processing of whole-brain MRSI. MRSIPrep accepts quantified metabolite maps and associated quality metrics as input and performs automated quality control, tissue correction, spatial normalization, atlas projection, and derivative generation within a standardized workflow. The framework produces harmonized outputs suitable for conventional regional or voxel-based analyzes and network-level applications.

MRSIPrep was designed according to four principles. *Reproducibility*: the framework is distributed exclusively as a Docker image bundling every external dependency (ANTs [20], FSL [21, 22], FreeSurfer [23], PETPVC [24], Chimera [25]) at fixed versions, so that a given dataset and configuration produce the same derivatives regardless of the host system. *Modularity*: is provided by independently configurable and cacheable Nipype [26] workflow components. *Transparency*: MRSIPrep generates an automated quality-control report for every processing run, summarizing spatial registration accuracy, metabolite coverage, voxel-and region-level quality metrics, tissue composition, and atlas projection, so that every stage can be visually inspected rather than trusted blindly. Finally, *analysis agnosticism* allows the same derivatives to support voxelwise, regional, and network-level analyses.

We evaluate whether these properties hold across heterogeneous whole-brain MRSI data. Because MRSIPrep operates downstream of spectral quantification, its processing is independent of the acquisition trajectory and fitting software. We therefore test this design across three whole-brain FID-MRSI implementations spanning high and ultra-high-field imaging conditions: 3D Cartesian CS-SENSE FID-MRSI [9] at 3 T, non-Cartesian ECCENTRIC FID-MRSI [5] at 3 T and 7 T, and concentric-ring trajectory FID-MRSI (CRT-FID-MRSI) [27] at 7 T. Registration and voxel-based-analysis performance are first assessed against synthetic ground truth (§3), after which the same pipeline is applied to clinical and healthy-control datasets spanning five acquisitions, three sites, two field strengths, and three MRSI sequences (§3.2).

## 2 Methods

### 2.1 Overview

MRSIPrep is implemented as a per-recording workflow built on Nipype [26], following the design pattern established by the NiPreps community [18, 19] for MRI/fMRI postprocessing: every stage is implemented as an independently cacheable node, so a partially completed or reparameterized run recomputes only the stages whose inputs or configuration changed rather than the recording from scratch. The pipeline is distributed exclusively as a Docker image bundling ANTs, FSL (FAST and FLIRT/FNIRT), FreeSurfer (recon-all, mri_synthseg), PETPVC, and Chimera, so that all processing runs against a fixed set of dependency versions regardless of the host system.

MRSIPrep takes as input quantified metabolite maps, associated voxelwise quality maps (Cramér–Rao lower bounds (CRLB), signal-to-noise ratio (SNR), linewidth/full-width-at-half-maximum (FWHM)), and a T1-weighted anatomical image, organized as a BIDS dataset. An optional dataset-level mrsinmrs.json sidecar, following the MRSinMRS minimum reporting standard [17], supplies acquisition parameters (e.g. repetition time, flip angle, field strength) that are surfaced directly in the quality-control report regardless of whether T1 saturation correction (§2.3.4, itself optional) is requested. Processing proceeds through two converging tracks (**Figure 1**): an anatomical track that segments and spatially normalizes the T1-weighted image, and an MRSI track that preprocesses and registers the metabolite maps to that anatomical reference. After the tracks converge at registration, MRSIPrep performs tissue correction, atlas projection, regional summarization, and optional metabolic connectivity estimation, concluding with an automated quality-control report; each stage is detailed in the corresponding subsection below. Each recording is processed by its own independent workflow instance, so multi-subject batches parallelize across recordings (–nproc) while the stages within one recording execute as a linear dependency chain. Every stage writes its derivatives as a BIDS-Derivatives dataset under derivatives/mrsiprep/, named following BIDS entity conventions (Table 1).

**Figure 1:**
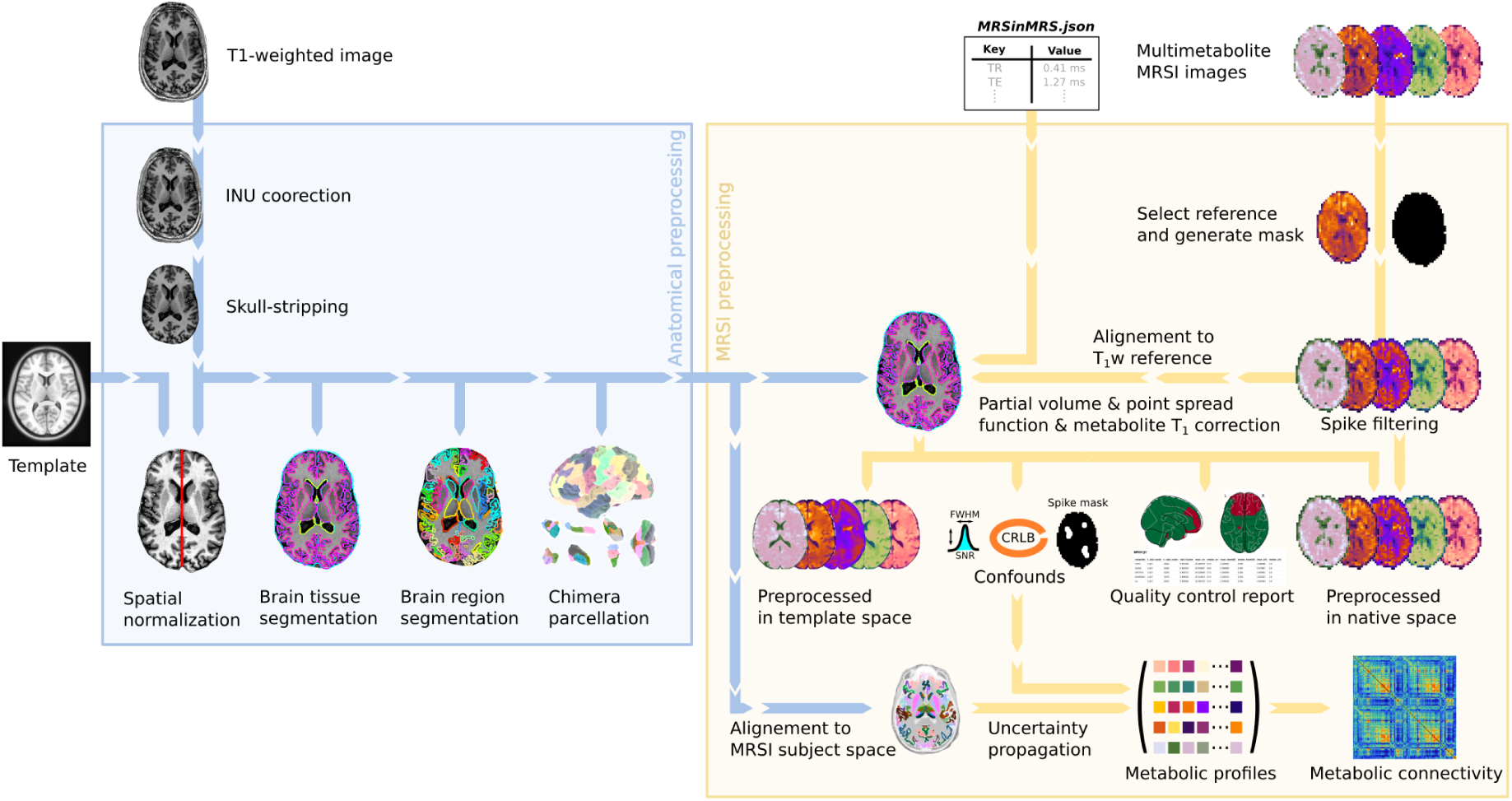
MRSIPrep workflow architecture. The anatomical track (blue) performs bias-field (INU) correction and skull-stripping on the T1-weighted image, then spatial normalization to a template, brain tissue segmentation, brain region segmentation, and Chimera multi-atlas parcellation. The MRSI track (yellow) selects a reference metabolite map and generates a brain mask, applies spike filtering, and registers the reference to the T1w image before partial-volume correction. The corrected maps are resampled into native and template space alongside voxelwise confound maps (FWHM, SNR, CRLB, spike mask) and an automated quality-control report. The Chimera parcellation is aligned into native MRSI subject space, where CRLB-based uncertainty propagation yields regional metabolic profiles and metabolic connectivity matrices.

**Table 1:**
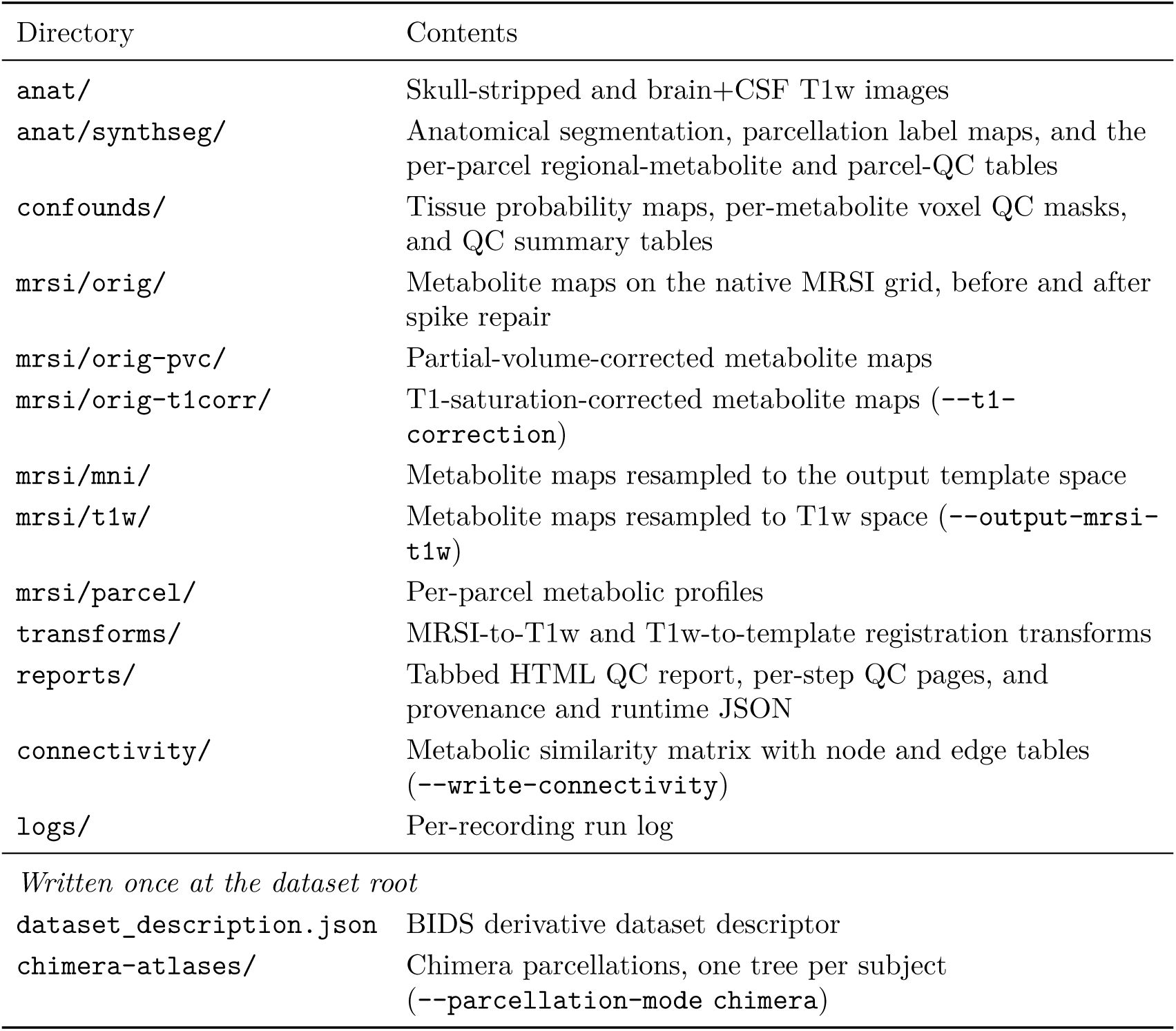
MRSIPrep output structure. Directories written per recording under sub-<LABEL>/ses-<LABEL>/, plus the entries written once at the dataset root. File names carry BIDS-style entities (space-, met-, atlas-, desc-) naming the output space, metabolite, atlas, and processing step. Rows marked with a flag are produced only when that flag is passed.

| Directory | Contents |
| --- | --- |
| `anat/` | Skull-stripped and brain+CSF T1w images |
| `anat/synthseg/` | Anatomical segmentation, parcellation label maps, and the per-parcel regional-metabolite and parcel-QC tables |
| `confounds/` | Tissue probability maps, per-metabolite voxel QC masks, and QC summary tables |
| `mrsi/orig/` | Metabolite maps on the native MRSI grid, before and after spike repair |
| `mrsi/orig-pvc/` | Partial-volume-corrected metabolite maps |
| `mrsi/orig-t1corr/` | T1-saturation-corrected metabolite maps (`--t1-correction`) |
| `mrsi/mni/` | Metabolite maps resampled to the output template space |
| `mrsi/t1w/` | Metabolite maps resampled to T1w space (`--output-mrsi-t1w`) |
| `mrsi/parcel/` | Per-parcel metabolic profiles |
| `transforms/` | MRSI-to-T1w and T1w-to-template registration transforms |
| `reports/` | Tabbed HTML QC report, per-step QC pages, and provenance and runtime JSON |
| `connectivity/` | Metabolic similarity matrix with node and edge tables (`--write-connectivity`) |
| `logs/` | Per-recording run log |
| *Written once at the dataset root* |  |
| `dataset_description.json` | BIDS derivative dataset descriptor |
| `chimera-atlases/` | Chimera parcellations, one tree per subject (`--parcellation-mode chimera`) |

### 2.2 Anatomical Preprocessing

The T1-weighted image is bias-field corrected and skull-stripped, then segmented with SynthSeg [28], a contrast-and resolution-agnostic deep-learning segmentation tool that requires no prior intensity normalization or template-specific tuning. SynthSeg-based brain extraction retains the whole brain brain mask comprising gray matter (GM), white matter (WM), ventricles, and inner/outer cerebrospinal fluid (CSF) and extra-ventricular CSF. The full CSF compartment remains available for tissue-fraction estimation with FSL FAST [29]. Two tissue-segmentation paths are available: a fast path (synthseg-fast, the default) using SynthSeg’s cortical parcellation directly, and a full path invoking FreeSurfer’s recon-all [23] when surface-based parcellation is required (–parcellation-mode chimera). In that mode, recon-all always precedes the Chimera step: Chimera [25, 30] consumes recon-all’s surface reconstruction and subcortical segmentation as input rather than deriving them itself, so MRSIPrep runs recon-all to completion first (skipped only if a valid FreeSurfer subject directory already exists for that subject) before invoking Chimera. Chimera then fuses independently sourced, region-specific atlases into a single whole-brain parcellation, one atlas per supra-region, selected by a 10-character scheme code identifying which source atlas fills each position, rather than adopting any single atlas’s own definition of every structure. The scheme code’s ten positions are, in order: cortex, subcortical/basal ganglia, thalamus, amygdala, hippocampus, hypothalamus, cerebellum, brainstem, gyral white matter, and white matter. MRSIPrep’s default scheme is LFMIHIFIFF; the scheme used in this paper’s results (§3.2, §2.3.9), LFMIHISIF (scale 3 in most analyses; scale 1 where noted), omits the gyral-white-matter position, leaving a 9-character code: **L**, the Lausanne multi-scale cortical parcellation [31], for cortex; **F**, FreeSurfer/Aseg whole-brain segmentation [32], for subcortex; **M**, MIALThalParc, for the thalamus [33]; **I**, FSAmygHippoParc, for the amygdala [34]; **H**, HBT, for hippocampal subfields [35]; **I**, FSHypoThalParc, for the hypothalamus [36]; **S**, SUIT, for the cerebellum [37]; and **I**, FSBrainStemParc, for the brainstem [38]. All parcellations are built in the subject’s own FreeSurfer native space before being carried through to MRSI-space projection (§2.3.7).

### 2.3 MRSI Preprocessing

#### 2.3.1 Reference Metabolite and MRSI Brain Mask

When present for a recording, the requested reference metabolite (–ref-met) is used directly; otherwise MRSIPrep falls back to the voxelwise mean across the available metabolites. The MRSI-space brain mask follows the same fallback logic: a pre-existing mask is used as-is; otherwise a thresholded water-reference map; otherwise the union of nonzero signal across all metabolite maps.

#### 2.3.2 Spike Filtering

Voxelwise spike filtering flags outlier voxels in every metabolite map (values above the 99th percentile of in-brain signal by default, –spikepc) and repairs them prior to any downstream step that would otherwise be biased by artifactual amplitudes: a local 3 × 3 × 3 median replacement, followed by biharmonic inpainting of any voxel still zero inside the brain mask, then smoothing (FWHM matched to the native MRSI voxel size by default) spliced back in only at the repaired locations rather than across the whole map.

A percentile threshold alone cannot separate real focal signal from acquisition noise, since an isolated single-voxel outlier and a genuine, spatially coherent abnormality can both exceed the same cutoff. The spike filtering step therefore exempts connected spike clusters larger than a maximum size (–spike-max-cluster-voxels) from repair, treating them as more likely to reflect real signal than artifact.

Calibrating these thresholds correctly matters beyond artifact removal: MRSI has been used clinically to detect and delineate tumor lesions precisely because abnormal tissue can carry metabolite levels that are large, multi-standard-deviation outliers relative to normal-appearing tissue. McKnight et al. [39] validated a Cho-to-NAA index, expressed in standard deviations from a control-voxel population, against histopathology and found tumor presence was best predicted above 2.5 standard deviations (SD) (90% sensitivity, 86% specificity); Zhong et al. [40] reported that a two-fold choline/NAA elevation over contralateral normal-appearing white matter already indicates substantial (∼30–35%) tumor infiltration within a voxel, with tumor cores reaching five-fold elevations. A spike filter’s cluster-size cap and z-score ceiling must leave some margin below these clinically established magnitudes. A too aggressive setting risks smoothing away exactly the kind of focal, elevated-amplitude signal that constitutes a genuine finding rather than an artifact.

#### 2.3.3 Confound Outputs

MRSIPrep exposes every quantity that encodes a preprocessing assumption as an explicit, per-metabolite confound map, written under a dedicated confounds/ output folder rather than folded silently into the metabolite values themselves or scattered alongside the signal maps they describe. Three categories are distinguished by provenance. *Inherited confounds* such as CRLB, SNR, and linewidth/FWHM which originate upstream, from the spectral-fitting step that produced the input metabolite maps, and are treated as read-only quality descriptors that MRSIPrep carries through registration, resampling, and parcellation unchanged (§2.3.9). *Injected confounds*, include the per-metabolite spike mask (§2.3) are instead generated by MRSIPrep’s own signal-processing decisions and would not exist without it. *Derived confounds* comprise the GM/WM/CSF tissue-fraction maps (§2.3.6), originate from the anatomical image via MRSIPrep’s own tissue-segmentation step, and describe the partial-volume composition of each MRSI voxel independent of any particular metabolite. All three categories are written as separate NIfTI volumes (one per metabolite and space, where applicable) since they are spatially varying quantities at MRSI resolution, distinguished by the same space/met/desc filename entities used throughout MRSIPrep’s outputs; per-metabolite spike masks are propagated into T1w/MNI space only when explicitly requested (–transform-spikemask), while CRLB, SNR, and FWHM maps are always transformed alongside the signal maps they describe. Making this distinction explicit lets downstream users judge whether a given voxel’s quality reflects genuine acquisition/fitting uncertainty, a correction MRSIPrep itself applied, or the voxel’s underlying tissue composition.

#### 2.3.4 T1 Saturation Correction

Raw fitted metabolite amplitudes may be systematically underestimated when the acquisition repetition time (TR) is short relative to a metabolite’s own longitudinal relaxation time (*T*_1_), since the spin system has not fully recovered between excitations. MRSIPrep does not correct for this by default (–t1-correction none); opting in (–t1-correction literature) applies a single scalar correction factor per metabolite and recording as a *protocol-level* correction, not a per-voxel one and derived from the steady-state spoiled-gradient-echo signal equation

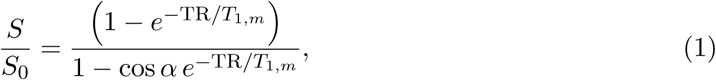

where *α* is the nominal flip angle and *T*_1*,m*_ is a literature value for metabolite *m* at the acquisition’s field strength; each metabolite map is multiplied by 1*/*(*S/S*_0_) to approximate its fully relaxed amplitude [41]. TR, flip angle, and field strength are read from a dataset-level mrsinmrs.json sidecar following the MRSinMRS reporting standard [17]; because that standard imposes no fixed field-naming schema, MRSIPrep validates a small whitelist of recognized key spellings and fails a recording outright rather than silently proceeding if a required field is absent or if two recognized spellings disagree.

Literature *T*_1_ values are curated in a data-only JSON configuration (editable independently of the pipeline source) keyed by metabolite and field strength, each entry recording the point estimate, its reported uncertainty, tissue/resonance context, source citation, and a curation status (*verified*, *proxy*, *derived*, or *todo*). Values are drawn from direct 3 T and 7 T metabolite relaxometry [42–44]. Lookup is exact: a metabolite is matched only against its own literature entry at the recording’s own field strength, with no fallback to a similarly named metabolite and no interpolation across field strengths; a *todo*-status or absent entry raises an explicit error identifying the gap. Because the input metabolite maps may already be water-referenced upstream by the quantification pipeline, –t1-correction-water-status records whether that water-scaling step has already applied its own *T*_1_ correction (corrected, uncorrected, or the conservative default unknown, which applies the metabolite-only correction and flags the ambiguity in the output provenance); this status cannot be inferred automatically, since MRSinMRS has no dedicated field for it.

T1 saturation is a scanner/sequence-level property of the raw fitted amplitude, independent of both the individual-voxel outlier repair of §2.3 and the spatial partial-volume redistribution of §2.3.6; MRSIPrep therefore applies it after spike filtering and before partial-volume correction, so that PVC’s own overshoot guard compares against the true, *T*_1_-recovered amplitude rather than the raw one. Corrected maps are written as a distinct derivative (desc-signalt1corr) alongside a per-recording confound table documenting each metabolite’s assumed *T*_1_, the acquisition parameters used, the resulting correction factor, and its sensitivity to the literature value’s reported uncertainty.

#### 2.3.5 Spatial Registration and Normalization

MRSI maps are registered to the T1-weighted reference with ANTs [20]. MRSIPrep supports two interchangeable registration backends, selected via –registration-backend: ANTs (antsRegistrationSyN.sh) and FSL (FLIRT linear registration [21], optionally followed by FNIRT [22] deformable refinement). For the ANTs backend, two transform models are available: rigid-plus-affine (-t a, “ANTs (R+Aff)”) and rigid-plus-deformable SyN (-t s, “ANTs (R+SyN)”), trading additional nonlinear correction for compute cost. Normalization to template space (triggered by –output-spaces MNI152NLin2009cAsym, MRSIPrep’s default output space, or by –parcellation-mode atlas, which requires it) follows the same backend choice and resamples every metabolite, CRLB, SNR, and FWHM map into the selected space at a configurable resolution, set per space with a resmodifier (–output-spaces MNI152NLin2009cAsym:res-2; MRSI-native, T1w-native, or an explicit isotropic value). For longitudinal datasets, –longitudinal instead builds one unbiased T1w subject-template with ANTs across sessions and registers that template to the target space, avoiding that each session registers independently to the target space. We benchmark all four backend/transform combinations directly against one another in §3.

#### 2.3.6 Partial-Volume Correction and Point Spread Function Correction

Because each MRSI voxel typically contains a mixture of tissue classes **Y** = {tissue_1_*, …,* tissue*_n_*}, the observed metabolite signal *M_v_* in voxel *v* can be modeled as a tissue-weighted mixture:

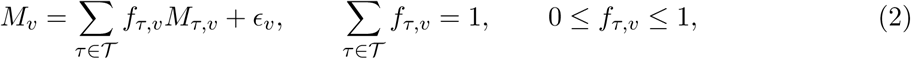

where *f_τ,v_* denotes the fraction of tissue class *τ* in voxel *v*, and *M_τ,v_* denotes the tissue-specific metabolite signal contribution. Here, we consider the tissue classes relevant to the MRSI signal, namely GM, WM, and CSF whose fractions are estimated using FAST and resampled into MRSI space. MRSIPrep solves this mixture problem with PETPVC’s region-based voxelwise (RBV) algorithm [24], applied to every metabolite map jointly with the stacked GM/WM/CSF fraction volume and an isotropic point-spread function (PSF) for deconvolution. Following PETPVC’s modeling convention, the PSF is approximated as a three-dimensional Gaussian kernel parameterized by its full width at half maximum (FWHM) along each axis. The PSF full-width defaults to the MRSI acquisition’s own native resolution (the mean voxel size of the reference metabolite map), since that is the true width of the spatial response function being deconvolved and it otherwise varies severalfold between protocols (e.g. 5.0–5.2 mm at 3 T vs. 3.4–3.5 mm at 7 T for the ECCENTRIC acquisitions in §3); an explicit override is also supported. Partial-volume correction is enabled by default and can be disabled with –no-pvc, in which case the uncorrected mixture in Eq. 2 is carried forward unmodified.

#### 2.3.7 Atlas Projection and Regional Derivatives

MRSIPrep supports two parcellation backends: Chimera [25] (§2.2), a multi-atlas fusion of independently sourced cortical, subcortical, thalamic, amygdalar, hippocampal, hypothalamic, cerebellar, brainstem, and white-matter atlases, and a bundled MNI-space atlas requiring no FreeSurfer license. For each metabolite *m* and atlas parcel *p*, MRSIPrep computes a per-parcel summary as an SNR-weighted average over quality-passing voxels:

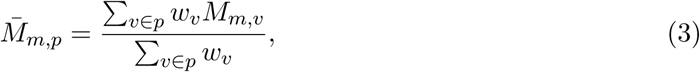

where *M_m,v_* is the amplitude of metabolite *m* in voxel *v*, the sum runs over voxels in parcel *p* that pass the metabolite’s QC mask, and *w_v_* is the voxel’s SNR (uniform weighting when no SNR map is available). Unweighted mean, median, standard deviation, coverage fraction, and mean CRLB/linewidth/tissue-fraction are reported alongside the weighted mean for every parcel.

#### 2.3.8 Regional Metabolic Profile Estimation and Uncertainty Propagation

Beyond the per-parcel summary table of §2.3.7, MRSIPrep always constructs a normalized *regional metabolic profile* for each retained parcel, independent of –parcellation-mode, a representation designed specifically to support inter-regional comparison, and the shared input to any downstream analysis that treats a parcel’s metabolite panel as a single feature vector (metabolic connectivity, §2.3.9; gradient mapping). Profile estimation is a standard output for every recording and does not require –write-connectivity. Construction begins with a voxel-wise z-score of each metabolite map *m* over the whole brain volume (subtracting the participant-and metabolite-specific whole-brain mean *µ_m_* and dividing by its standard deviation *σ_m_*), so that no single metabolite’s concentration scale dominates the profile:

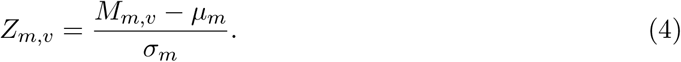

The z-scored map is then averaged within each retained parcel *p* as a gray-matter-fraction-weighted mean over quality-passing voxels,

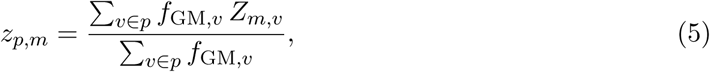

giving a *K*-dimensional regional metabolic profile **m***_p_* = [ *z_p,_*_1_*, z_p,_*_2_*, …, z_p,K_*] for the *K* metabolites in –metabolites. Parcels with poor CRLB or insufficient coverage are flagged in the quality-control report (§2.4) using the same regional QC fields as §2.3.7 (coverage fraction, mean CRLB/linewidth), leaving the decision of whether to exclude them to the user.

MRSIPrep uses the Monte Carlo uncertainty-propagation approach of Instrella and Juchem [45]: for each metabolite map, every voxel’s signal *M_m,v_* is perturbed *K*_pert_ times (default *K*_pert_ = 50, –connectivity-n-perturbations) according to its CRLB-derived variance by drawing

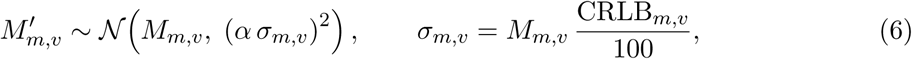

where CRLB*_m,v_* is the voxel’s Cramér–Rao lower bound (expressed as a percentage, as reported by the spectral-fitting tool) and *α* is a fixed noise-scaling factor (default *α* = 2, –connectivity-sigma-scale). Perturbed values are clipped to 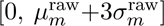 (pre-perturbation map mean and standard deviation) to suppress non-physiological outliers before each perturbed volume is carried through the same voxelwise z-score (Eq. 4) and parcel averaging (Eq. 5) as the original map. Stacking the original profile with its *K*_pert_ perturbed counterparts yields an augmented per-parcel feature matrix **X***_p,_*: ∈ R*^K^*^(*K*pert+1)^, propagating per-voxel quantification uncertainty into every downstream use of the profile rather than treating each region’s metabolite level as noise-free.

#### 2.3.9 Metabolic Connectivity

We use the term *metabolic connectivity* to denote an edge-wise relationship between brain regions obtained by comparing their representations over the metabolic signal domain encoded in the chemical-shift dimension measured by MRSI. This parallels functional connectivity, in which regional BOLD signals are compared over the temporal domain, and structural connectivity, in which anatomical connection evidence is inferred from diffusion information distributed along physical white-matter trajectories.

For MRSI, such an edge could in principle be defined directly from regional spectra over the chemical-shift domain. Raw spectra, however, remain sensitive to noise, baseline distortions, linewidth variation, and sequence-dependent spectral characteristics. MRSIPrep therefore constructs connectivity from the post-quantification regional profiles described in §2.3.8. These profiles provide a lower-dimensional and biochemically interpretable representation of the spectral domain, in which each region is characterized by its pattern of quantified metabolites and derived profile features.

Both the parcel-wise summaries of §2.3.7 and the regional profiles of §2.3.8 are generated unconditionally for every recording, independent of –parcellation-mode. Connectivity-matrix construction, by contrast, is an explicit opt-in operation (–write-connectivity) applied to these already-computed profiles rather than a default preprocessing output. This design follows the broader convention established by fMRIPrep [18], which produces preprocessed BOLD data and associated confounds while leaving functional-connectivity estimation to downstream analysis tools such as XCP-D [46].

MRSIPrep nevertheless provides a reference connectivity implementation, generalizing the procedure used to construct the metabolic connectome in Lucchetti et al. [8] to arbitrary atlases and metabolite panels. Users may instead construct alternative connectivity estimates directly from the exported regional profiles using any suitable similarity or dependence measure. Given the augmented profile matrix **X** ∈ R*^N^*^×*K*(*K*pert+1)^ defined in §2.3.8, where *N* is the number of retained parcels, the metabolic similarity between parcels *i* and *j*, forming one entry of the metabolic connectivty matrix, is

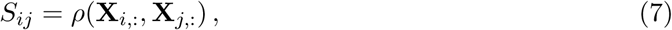

where *ρ* denotes a configurable similarity function (–connectivity-method). The default is Spearman correlation, as used in Lucchetti et al. [8].

### 2.4 Quality-Control Framework

MRSIPrep summarizes quality at the voxel, regional, and subject levels. Voxel inclusion is governed by metabolite-specific criteria on linewidth, SNR, CRLB, tissue composition, and spatial coverage. An automated per-recording HTML report summarizes spatial registration accuracy (with the reference-metabolite overlay always generated regardless of whether T1w-space MRSI maps are retained as permanent derivatives), metabolite coverage, voxelwise quality metrics, tissue composition, and atlas projection, so that every processing stage can be visually inspected without re-running the pipeline. When an optional mrsinmrs.json file is present at the BIDS dataset root, the report additionally renders a dedicated MRSinMRS section [17] listing the recording’s sequence, hardware, and reconstruction parameters alongside a citation block, so that a study’s acquisition provenance and its processing provenance are shared in a single, versionable artifact rather than documented separately. Published acquisition parameter sets can be reproduced directly via –config-preset, with the report crediting the originating study. A full example report, rendered end-to-end (Chimera parcellation, metabolic connectivity) for a subject from the public SynthMRSI-Project dataset (§2.6), is available at https://github.com/MRSI-Psychosis-UP/MRSIPrep/tree/main/docs/examples/sub-01_ses-01.

### 2.5 Statistical Analysis

Regional profile replicability (§3.2) is assessed via Spearman correlation between the two cohorts’ region-level mean profiles, each cohort’s per-region value being the median across its subjects. Because root regions are not independent, exchangeable units under a plain parametric test (neighboring regions share spatial autocorrelation from biology, partial-volume correction, registration smoothing, and gray-matter gradients, which anti-conservatively inflates naive Spearman significance), significance is instead assessed against a spatially-aware null: variogram-matched surrogate maps [47] (brainsmash), built from a Euclidean distance matrix between root-region centroids of the bundled chimera-LFMIHISIF-1 atlas. Because this parcellation spans cortex, subcortex, thalamus, cerebellum, and hippocampus rather than a single continuous cortical sheet, a classical spherical spin test does not apply. For each metabolite, 5,000 surrogates are generated independently for each side of the comparison (preserving that side’s own spatial autocorrelation while destroying its true correspondence with the other side); the reported *p*-value is the more conservative (larger) of the two resulting one-sided empirical tests.

### 2.6 Validation Datasets

Two dataset families support the validation reported below. **Real acquisitions**: two single-subject recordings, each with a matched MP2RAGE anatomical, used for the registration-accuracy benchmark (§3.1.1) and the Supplementary Material’s runtime-scaling benchmark: **Geneva3T** (3D Cartesian CS-SENSE FID-MRSI [9]; Siemens MAGNETOM TrioTim, 32-channel coil, 5.0 × 5.0 × 5.2 mm^3^ nominal MRSI resolution) and **Geneva7T** (3D ECCENTRIC FID-MRSI [5]; Siemens MAGNETOM Terra.X, 32-channel coil, 3.4 × 3.4 × 3.5 mm^3^).

**Synthetic ground truth**: a 32-subject synthetic validation cohort built by warping model-synthesized MRSI metabolite signal, with a deterministic region-specific abnormality injected into half the subjects – CrPCr into bilateral Precuneus (AAL region IDs 67/68) and GluGln into bilateral Thalamus (AAL region IDs 77/78) – onto real T1w anatomicals, used for the detection-validation benchmark (§3.3), where the true injected region provides an exact ground truth independent of registration backend. The injected abnormality’s metabolite, region, and magnitude are an arbitrary methodological choice, not a modeled pathology or literature-derived effect size: CrPCr/Precuneus and GluGln/Thalamus were chosen only for their anatomical separation and non-overlapping tissue composition, and no biological or clinical significance should be attached to this regional metabolite pairing. Its sole purpose is to create a spatially localized, ground-truth-known signal against which detection sensitivity can be compared across backends. For CrPCr specifically, ground truth is available in two forms: the raw AAL Precuneus parcel described above, and a gray-matter-precise version restricted to SynthSeg’s own DKT cortical Precuneus labels (inherently gray-matter-only, unlike the AAL parcel’s ∼62% gray-matter native-space composition), used for a stricter boundary-tracking follow-up in the same section. The model-synthesized signal itself comes from a 3D U-Net trained to predict per-voxel signal for the same 5 metabolite channels from a T1-weighted image alone, using real template subjects’ T1w and metabolite maps warped onto a shared MNI grid; at inference, the trained network produces synthetic metabolite signal in that same MNI space for a new T1w input, and the inverse of a randomly sampled, precomputed real subject’s own MRSI-to-MNI transform is then applied to resample this synthetic signal into that subject’s native MRSI resolution and geometry, so each dummy subject carries a real acquisition’s native-space grid and that subject’s own empirical CRLB/SNR/FWHM quality maps rather than predicted ones. A related public, CC0-licensed synthetic dataset (32 subjects, real T1w with model-synthesized MRSI signal but no group-level injection) is available at https://doi.org/10.5281/zenodo.21477047 for independent testing of the pipeline itself.

**Independent multi-site datasets**: real-world healthy-control cohorts spanning five acquisitions across three sites, two field strengths, and three MRSI sequence implementations, used for the cross-site/cross-sequence regional profile replicability check (§3.2). Acquisition parameters for all five are summarized in **Supplementary Table 5**, following the MRSinMRS minimum reporting standard [17].

**Lausanne3T-FID** (*n* = 12) and **Lausanne3T-ECCENTRIC** (*n* = 15) are two sequences acquired on the same Lausanne 3 T scanner in healthy participants recruited alongside the Lausanne psychosis cohort [48]; participants were required to be free of any lifetime psychiatric diagnosis and to have no first-degree relative with a psychosis-related disorder. The study was approved by the Cantonal Ethics Committee for Research on Human Beings, Vaud (CER-VD PB_2017-00675 and 2024-01777), and all participants provided written informed consent.

**Geneva3T-FID** (*n* = 59), the same Cartesian CS-SENSE sequence as Lausanne3T-FID acquired on an independent 3 T scanner in Geneva, is drawn from the Mindfulteen study [49]. Participants were aged 13-15 years and were excluded for chronic somatic disease, significant medical conditions, psychotherapy in the previous 6 months, psychotropic medication in the previous month, or a current or past psychiatric disorder (except anxiety disorder or past major depressive disorder). The study was approved by the Geneva Cantonal Commission for Research Ethics (CCER 2018-01731), with written informed consent from a parent/legal guardian and assent from each participant.

**Geneva7T-ECCENTRIC** (*n* = 26) is drawn from an ongoing longitudinal study of the 22q11.2 deletion syndrome (22q11DS) in Geneva; participants younger than 13 years were excluded to better match the age range of the Lausanne and Geneva 3 T cohorts above. As with the other clinical cohorts, the study was approved by the competent cantonal ethics committee ((CCER 2020-02296), and participants and/or their legal guardians provided written informed consent.

**Vienna7T-FID** (*n* = 5) [27] is an independent, publicly available 7 T site and sequence, a concentric-ring-trajectory FID-MRSI (CRT-FID-MRSI) implementation distinct from both ECCENTRIC and Cartesian CS-SENSE. Its quantification pipeline resolves 9 metabolites as individually-resolved channels (CrPCr, GPCPCh, Ins, NAA, NAAG, Glu, Gln, GABA, GSH) rather than MRSIPrep’s usual 5 aggregate channels, since it never produces joint values such as tNAA for NAA+NAAG; for the cross-sequence comparison of §3.2, this restricts the shared parcellation to the 119 GM regions common to both the Vienna7T-FID and Geneva7T-ECCENTRIC scale-1 chimeraLFMIHISIF atlas (excluding brainstem sub-parcels: midbrain, pons, medulla, superior cerebellar peduncle). Vienna7T-FID’s quantification pipeline also did not export a per-metabolite CRLB map; rather than excluding it from validation, its regional profiles use MRSIPrep’s CRLB-fallback path (§2.3.8): a metabolite with no CRLB map is treated as 0% uncertainty (no injected noise), collapsing the usual CRLB-perturbed Monte Carlo draw to a single deterministic point estimate at the native MRSI signal value. For all four clinical/healthy-control cohorts above, and for participants with multiple scanning sessions, the first session with adequate whole-brain coverage was selected for analysis.

## 3 Results

### 3.1 Technical Validation

We first evaluate MRSIPrep’s registration accuracy on the two real ECCENTRIC acquisitions of §2.6 (runtime scaling on the same two acquisitions is reported in the Supplementary Material) and validate its voxel-based detection performance on the synthetic ground-truth cohort.

#### 3.1.1 Registration Accuracy Across Backends

–registration-backend offers ANTs (rigid+SyN for MRSI→T1w, rigid+affine+SyN for T1w→MNI by default) and FSL (FLIRT affine, with an optional FNIRT deformable stage). We compared four MRSI→T1w registration configurations, ANTs (Rigid+SyN), ANTs (Rigid+Affine), FSL FLIRT-only, and FSL FLIRT+FNIRT, against two T1w registration targets (brain-only and brain+CSF), on both real subjects (4 configurations × 2 targets × 2 subjects = 16 runs). Registration accuracy is reported as *signal-weighted leakage*: at each quality-passing voxel (resampled CRLB ≤ 20) the resampled reference-metabolite (CrPCr) signal magnitude is summed, and the leakage percentage is the fraction of that total signal mass falling outside the reference brain mask, deliberately not a raw voxel count, since a thin near-zero resampling artifact and a voxel carrying substantial misplaced signal would otherwise count identically.

**ANTs (Rigid+SyN), MRSIPrep’s default, has the least signal-weighted leakage at both field strengths** (Figure 2; 0.34% at 3 T, 0.44% at 7 T), roughly 6–10× less than FSL FLIRT-only (2.1%, 0.97%) and 12–16× less than FSL FLIRT+FNIRT (5.4%, 2.7%), consistent with the sharpest, most anatomically detailed overlay in Figure 2a. **ANTs (Rigid+Affine) is, unexpectedly, the second-leakiest configuration overall** (worse than either FSL variant at 7 T: 4.30% at 3 T, 4.73% at 7 T, roughly 11–12× the default ANTs pipeline), suggesting that adding a real affine correction on top of rigid alignment without a subsequent deformable stage lets the affine’s extra degrees of freedom fit noise in the relatively low-resolution, low-contrast MRSI reference rather than genuine anatomical correspondence. **FSL FLIRT+FNIRT leaks more than FLIRT-only at both field strengths** (5.4% vs. 2.1% at 3 T; 2.7% vs. 0.97% at 7 T), indicating that its deformable stage does not reduce real signal leakage despite its added runtime cost (11.6 min vs. 3.1 min at 3 T; 53.0 vs. 11.2 min at 7 T, nearly 2× ANTs (Rigid+SyN) itself at 7 T). Brain-only vs. brain+CSF as the T1w registration target has a small effect (≤0.5 percentage points) that flips direction between field strengths and is dominated by the choice of backend. Overall, ANTs (Rigid+SyN) remains the best combination of accuracy and runtime among the full-pipeline configurations tested.

**Figure 2:**
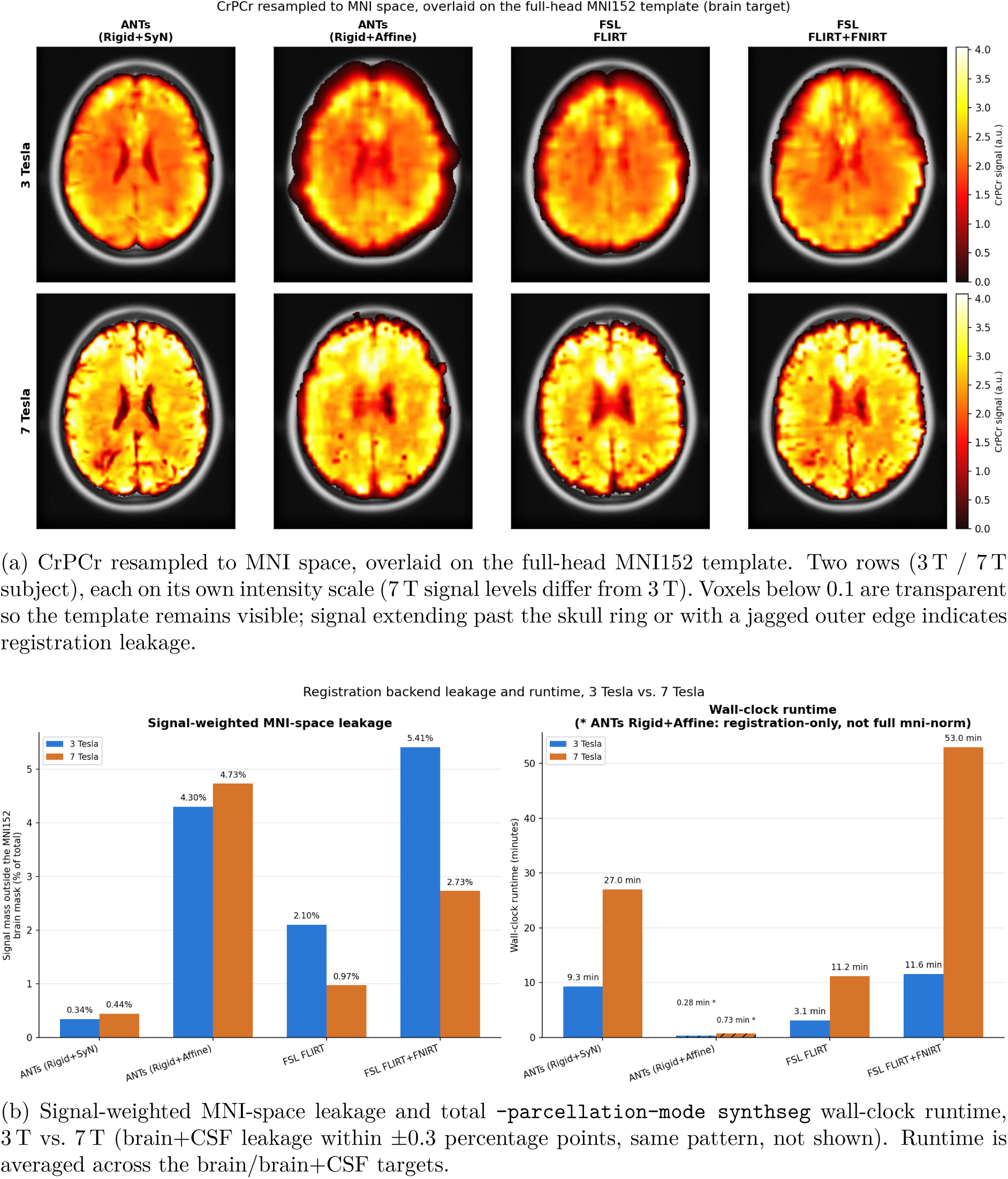
**Registration backend comparison in MNI space**, brain registration target, for the four configurations compared throughout (ANTs Rigid+SyN / ANTs Rigid+Affine / FSL FLIRT / FSL FLIRT+FNIRT, one column each in **(a)**, one bar group each in **(b)**). **(a)** shows the qualitative result and **(b)** quantifies it.

### 3.2 Cross-Site/Cross-Sequence Regional Profile Replicability

The benchmarks above validate MRSIPrep’s registration and detection performance on data it was built and tuned against; both are reproducibility evidence, since the same pipeline and configuration are applied to inputs it was developed against. A complementary test is MRSIPrep-derived regional metabolite profiles replicability on independent real-world data: different subjects, different scanners, different sites, and different MRSI sequences, processed with the same pipeline and parcellation (chimeraLFMIHISIF scale 1). Because metabolite units are not expected to agree across different sequences and field strengths (scanner-, sequence-, and reconstruction-dependent scaling and referencing choices affect the overall signal level independently of true tissue concentration), this is deliberately framed as a test of whether the relative spatial pattern of regional metabolite levels replicates across acquisitions. Three comparisons are reported: two at 3 T, isolating in turn the effect of sequence (same site, different sequence) and of site (different site, same sequence), and one at 7 T. **Lausanne3T-FID vs. Lausanne3T-ECCENTRIC** (27 of 28 available recordings, one excluded for flagged spectral quality; *n* = 12 and *n* = 15) compares two sequences acquired on the same Lausanne scanner. **Lausanne3T-FID vs. Geneva3T-FID** (*n* = 12 and *n* = 59) compares the same Cartesian CS-SENSE sequence acquired on two independent 3 T scanners with slightly different repetition and echo times (see MRSinMRS parameters in Table 5). Both 3 T comparisons use the same 5 metabolites (CrPCr, GluGln, GPCPCh, NAANAAG, Ins) and the same 82-region chimeraLFMIHISIF scale-1 parcellation, GM-filtered (excluding only brain-stem-midbrain); each subject’s regional value is the median across the *K*_pert_ = 50 CRLB-perturbed draws of §2.3.8. **Vienna7T-FID vs. Geneva7T-ECCENTRIC** (*n* = 5 and *n* = 26) tests the same replicability question at 7 T, on the 9 metabolites Vienna7T-FID resolves individually and the 119-region parcellation this requires (§2.6), and using Vienna7T-FID’s CRLB-fallback profiles (§2.6); the other four datasets’ profiles all use the standard CRLB-perturbed estimation, reduced to a comparable per-subject point estimate via the median across draws. All three comparisons are assessed following §2.5.

**Regional profiles replicate strongly across every comparison and every metabolite tested (Table 2):** Spearman *ρ* = 0.85–0.93 for Lausanne3T-FID vs. Lausanne3T-ECCENTRIC, *ρ* = 0.63–0.83 for Lausanne3T-FID vs. Geneva3T-FID, and *ρ* = 0.68–0.90 for Vienna7T-FID vs. Geneva7T-ECCENTRIC, all significant under the spatially-aware null model (*p <* 2 × 10^−4^ for every metabolite except 7 T’s Gln and GSH, at *p* = 1 × 10^−3^ and 6 × 10^−4^ respectively). Absolute signal levels differ by orders of magnitude between acquisitions, but each dataset’s region-to-region pattern still matches closely once considered on its own scale. Lausanne3T-FID vs. Geneva3T-FID’s correlations are consistently lower than Lausanne3T-FID vs. Lausanne3T-ECCENTRIC’s (*ρ* = 0.63–0.83 vs. *ρ* = 0.85–0.93), despite comparing the same sequence. Together, the three comparisons span three independent sites, five acquisitions, and two field strengths. Full per-region trace plots for all three comparisons are provided in Supplementary Figures S1–S4 (Figures 7–10).

**Table 2:** Regional profile replicability across all three comparisons. *p*-values are empirical, from the spatially-aware null model of §2.5 (5,000 surrogates per side); *<* 2 × 10^−4^ indicates no surrogate in either direction ever matched or exceeded the observed correlation. The 3 T comparisons use 82 GM root regions; the 7 T comparison uses 119. Metabolites not fit as an individually-resolved channel in a given comparison are marked “–”. Full per-region trace plots are shown in Supplementary Figures 7–10.

| Metabolite | Lausanne3T-FID vs. Lausanne3T-ECCENTRIC |  | Lausanne3T-FID vs. Geneva3T-FID |  | Vienna7T-FID vs. Geneva7T-ECCENTRIC |  |
| --- | --- | --- | --- | --- | --- | --- |
| | Spearman $\rho$ | $p$-value | Spearman $\rho$ | $p$-value | Spearman $\rho$ | $p$-value |
| CrPCr | 0.90 | $< 2 \times 10^{-4}$ | 0.70 | $< 2 \times 10^{-4}$ | 0.90 | $< 2 \times 10^{-4}$ |
| GPCPCh | 0.93 | $< 2 \times 10^{-4}$ | 0.83 | $< 2 \times 10^{-4}$ | 0.90 | $< 2 \times 10^{-4}$ |
| Ins | 0.88 | $< 2 \times 10^{-4}$ | 0.64 | $< 2 \times 10^{-4}$ | 0.90 | $< 2 \times 10^{-4}$ |
| GluGln | 0.85 | $< 2 \times 10^{-4}$ | 0.63 | $< 2 \times 10^{-4}$ | – | – |
| NAANAAG | 0.89 | $< 2 \times 10^{-4}$ | 0.75 | $< 2 \times 10^{-4}$ | – | – |
| NAA | – | – | – | – | 0.88 | $< 2 \times 10^{-4}$ |
| NAAG | – | – | – | – | 0.82 | $< 2 \times 10^{-4}$ |
| Glu | – | – | – | – | 0.90 | $< 2 \times 10^{-4}$ |
| Gln | – | – | – | – | 0.68 | $1 \times 10^{-3}$ |
| GABA | – | – | – | – | 0.81 | $< 2 \times 10^{-4}$ |
| GSH | – | – | – | – | 0.72 | $6 \times 10^{-4}$ |

### 3.3 Quality-Control Performance

Beyond registration accuracy in the absolute sense above, we validate whether MRSIPrep’s pipeline (across the same four registration configurations) can recover the known, deliberately injected, and biologically arbitrary metabolic abnormality of §2.6 (CrPCr into Precuneus, GluGln into Thalamus) through a standard voxel-based-analysis (VBA) group comparison (FSL’s randomise -T, TFCE-corrected permutation test, 500 permutations, two-sample unpaired design). The injection region for each metabolite is shown directly as the blue ground-truth outline in **Figure 3**.

**Figure 3:**
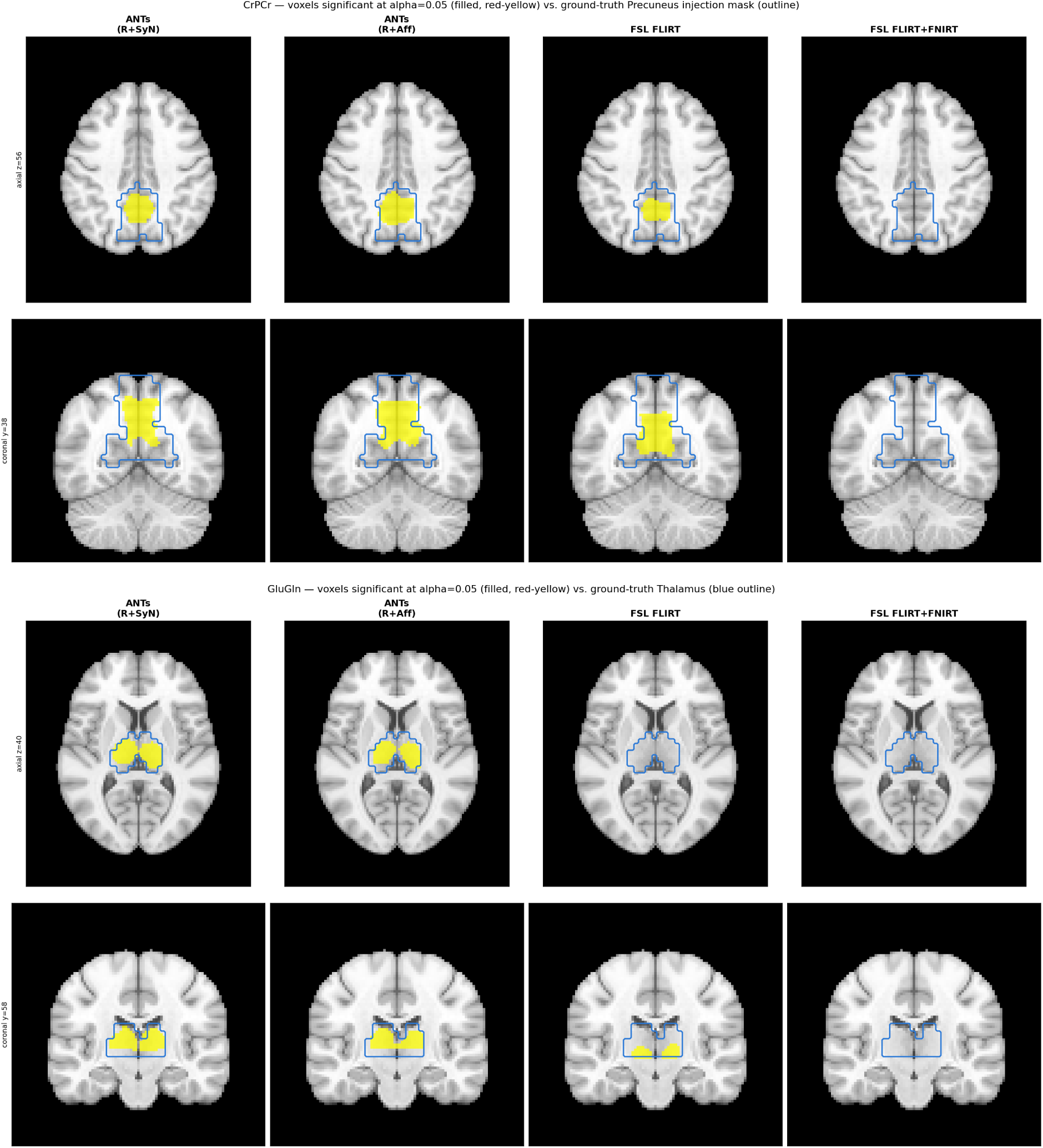
**CrPCr and GluGln detection:** voxels significant at *α* = 0.05 (filled) vs. the population ground-truth injection mask (blue outline), for all four backends. *Top (CrPCr):* ANTs (both configurations) and FSL FLIRT-only detect a cluster inside the true Precuneus region; FSL FLIRT+FNIRT detects no significant voxels at this slice. *Bottom (GluGln):* ANTs (both configurations) detects a clean bilateral cluster inside the Thalamus; FSL FLIRT-only detects a real but asymmetric cluster; FSL FLIRT+FNIRT detects zero significant voxels.

ROC and precision-recall are not redundant here: because the true injection region is a small fraction of total brain voxels, a backend can score near-chance on ROC-AUC while still showing high precision at very low recall (a handful of confident true positives), which ROC-AUC alone would not reveal. GluGln makes this concrete below: FSL FLIRT+FNIRT’s ROC-AUC (0.48, indistinguishable from chance) does not by itself indicate whether its corrp map carries any usable signal at all, whereas its PR-AUC (0.00) confirms it does not, ruling out the ROC-only reading that a small number of correct detections might still be recoverable at a stricter threshold.

**Figure 4:**
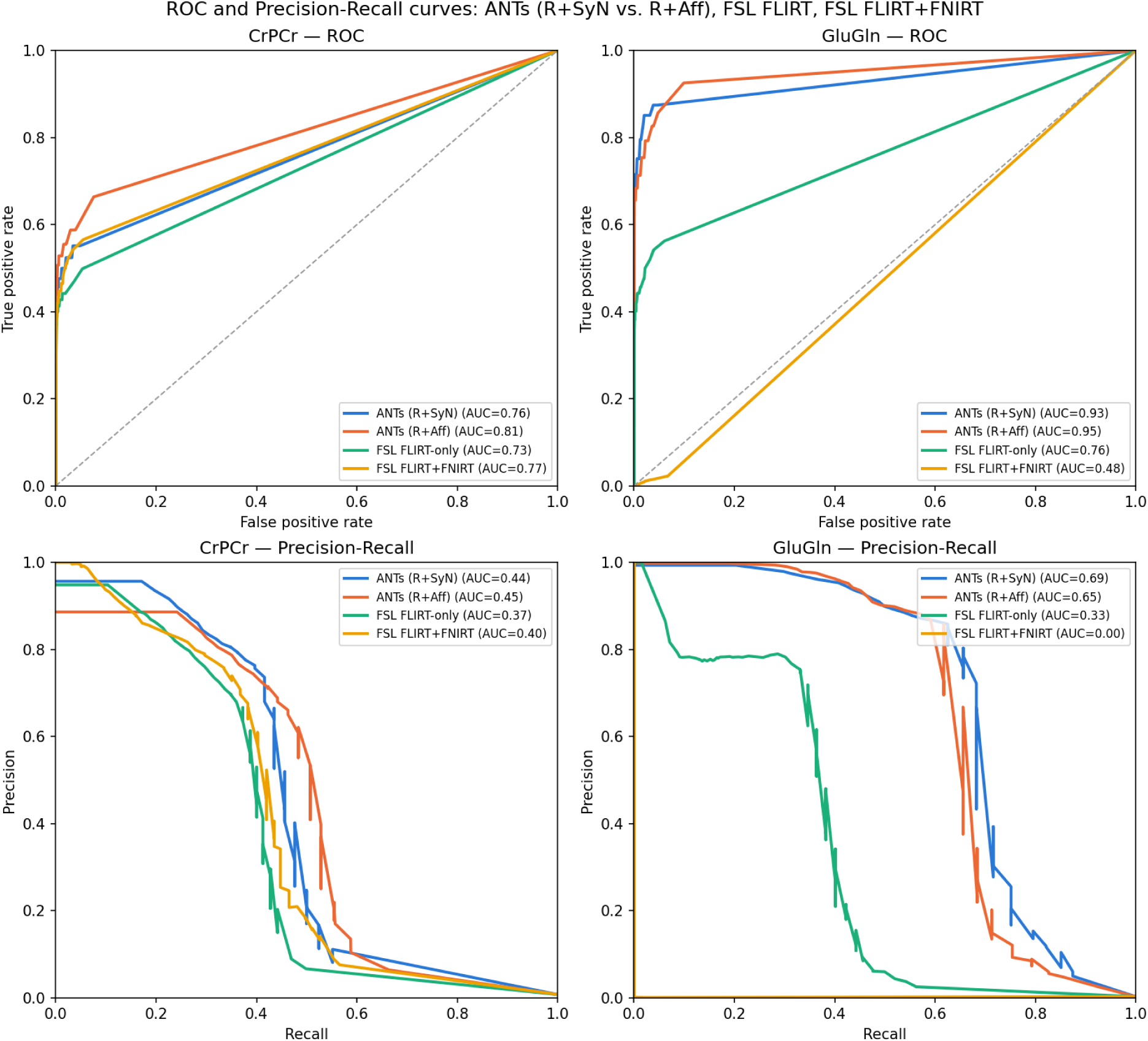
**ROC and precision-recall curves for CrPCr and GluGln**, sweeping the TFCE-corrected significance threshold across its full range, AUC in each legend.

**Table 3:** Threshold-independent detection performance by backend.

| Backend | CrPCr ROC-AUC | CrPCr PR-AUC | GluGln ROC-AUC | GluGln PR-AUC |
| --- | --- | --- | --- | --- |
| ANTs (R+SyN) | 0.76 | 0.44 | 0.93 | 0.69 |
| ANTs (R+Aff) | 0.81 | 0.45 | 0.95 | 0.65 |
| FSL FLIRT-only | 0.73 | 0.37 | 0.76 | 0.33 |
| FSL FLIRT+FNIRT | 0.77 | 0.40 | 0.48 | 0.00 |

**ANTs is the best-performing backend on both metabolites, and the R+Aff variant is consistently at least as good as the full R+SyN pipeline:** on GluGln it has the highest ROC-AUC of any backend (0.95), and on CrPCr it leads on both ROC-AUC (0.81) and PR-AUC (0.45); the deformable SyN stage does not clearly improve detection power over R+Aff on either metabolite. **FSL FLIRT+FNIRT is competitive with ANTs on CrPCr but collapses to near-chance on GluGln** (ROC-AUC 0.48, PR-AUC 0.00), consistent with §3.1.1’s finding that FNIRT’s nonlinear warp has higher signal-weighted leakage than ANTs or FLIRT-only. A small, deep, centrally-located structure like the Thalamus is exactly where local nonlinear-warp distortion does the most damage to a focal signal. FSL FLIRT-only is consistently the weakest backend on both metabolites but remains clearly above chance.

#### Gray-matter-precise boundary tracking

Bulk-overlap metrics like Dice cannot distinguish a detected cluster that tracks the true, convoluted gray-matter boundary from one that merely overlaps the same general neighborhood; the raw AAL Precuneus parcel used above is only ∼62% gray matter in native T1w space and sweeps into adjacent white matter rather than tightly tracing the cortical ribbon. This follow-up therefore uses the gray-matter-precise ground truth of §2.6 instead: CrPCr’s abnormality re-filtered and re-resampled through each backend’s already-computed registration transforms (no registration rerun, since registration depends only on anatomy, not the injected signal), scored with a symmetric mean surface distance and Hausdorff distance (mm) between the detected and ground-truth boundaries.

**Figure 5:**
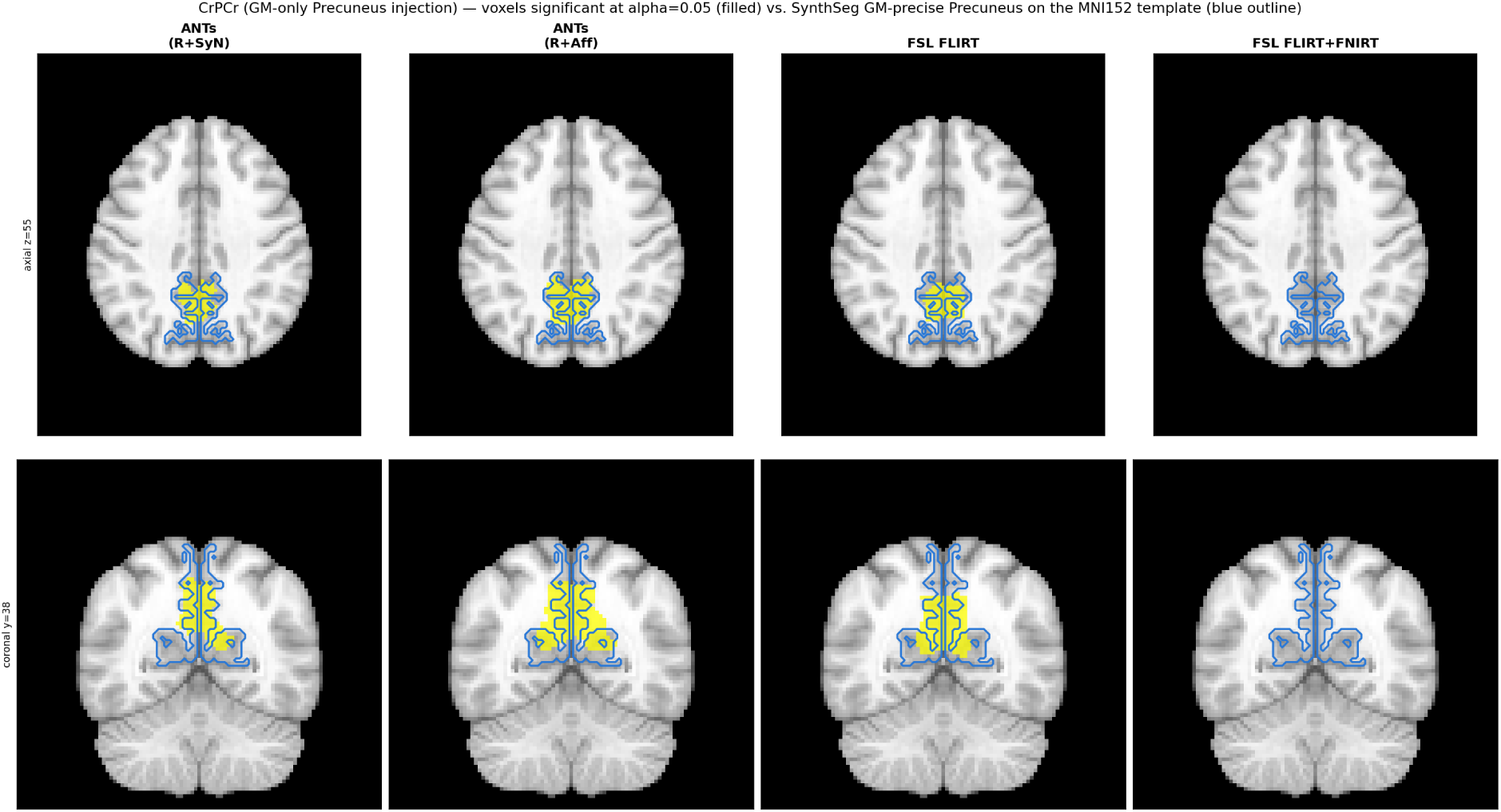
**CrPCr (gray-matter-only Precuneus injection) detection** vs. SynthSeg’s gray-matter-precise Precuneus segmentation of the MNI152 template itself (blue outline), all four backends.

**Table 4:** Gray-matter-precise Precuneus detection and boundary tracking.

| Backend | Dice | Sens. | Prec. | ROC-AUC | PR-AUC | Mean surf. dist. (mm) | Hausdorff (mm) |
| --- | --- | --- | --- | --- | --- | --- | --- |
| ANTs (R+SyN) | 0.341 | 0.244 | 0.569 | 0.810 | 0.315 | 5.47 | 23.07 |
| ANTs (R+Aff) | 0.432 | 0.374 | 0.510 | 0.849 | 0.326 | 4.32 | 22.36 |
| FSL FLIRT-only | 0.326 | 0.250 | 0.470 | 0.773 | 0.238 | 6.46 | 33.11 |
| FSL FLIRT+FNIRT | 0.017 | 0.009 | 0.721 | 0.815 | 0.268 | 17.94 | 45.52 |

**ANTs (R+Aff) has the best Dice, sensitivity, ROC-AUC, and boundary tracking of all four backends** on this harder, gray-matter-precise target (4.32 mm mean surface distance, under two and a half voxels at this 2 mm resolution); the deformable SyN stage reduces Dice here (0.341 vs. 0.432) relative to skipping it, mirroring the coarser AAL-parcel result above. **FSL FLIRT+FNIRT’s Dice collapses to 0.017** despite a ROC-AUC (0.815) close to ANTs SyN’s, and its boundary distance is roughly 3–4× farther from the true gray-matter boundary than either ANTs variant (17.94 mm mean, 45.52 mm Hausdorff, vs. ANTs (R+Aff)’s 4.32 mm/22.36 mm); its corrp map retains real discriminative signal, but that signal is spread too diffusely, or displaced, to ever cross the TFCE-corrected significance threshold in the right place. Across both the AAL-parcel and gray-matter-precise versions of this benchmark, ANTs’ deformable SyN stage never clearly outperforms the no-SyN configuration for this focal, planted-signal detection task.

### 3.4 Downstream Applications

MRSIPrep’s regional profile and connectivity modules (§2.3.8, §2.3.9) combine two mechanisms specifically intended to make connectomics-ready outputs more robust to known sources of MRSI-specific error, rather than treating every voxel’s metabolite value as an equally trustworthy point estimate. **CRLB-based uncertainty propagation**: rather than building a connectivity matrix from a single, noise-free regional value per parcel, each metabolite map is perturbed *K*_pert_ times according to its own voxelwise CRLB-derived variance (Eq. 6) before parcellation, so that parcels whose underlying voxels had less confident spectral fits contribute proportionally more variable, and therefore appropriately less overconfident. The same Monte Carlo approach was used by Instrella and Juchem [45] for uncertainty propagation in absolute MRS quantification. **Gray-matter-fraction weighting**: because each MRSI voxel is a partial-volume mixture of tissue types, and PETPVC’s RBV correction (§2.3.6) is applied at the signal level. The regional profile additionally weights each voxel’s contribution by its GM partial-volume fraction (Eq. 5) when tissue segmentation is available, so that voxels still dominated by white matter or CSF after PVC contribute less to a nominally gray-matter parcel’s profile.

Both mechanisms are methodologically motivated extensions of standard regional-profile construction, grounded in properties of the input data MRSIPrep already has available (per-voxel CRLB, per-voxel tissue fraction) rather than being specific to any one dataset; we do not report a quantitative comparison of connectivity-matrix stability with these mechanisms enabled versus disabled here. The improved robustness this CRLB-based uncertainty propagation confers on the resulting metabolic connectome, relative to a naive single-point-estimate construction, has already been demonstrated quantitatively by Lucchetti et al. [8].

As a downstream illustration, **Figure 6** shows the group-average metabolic connectivity matrix MRSIPrep produces for each of the five real-world datasets introduced in §3.2 (Lausanne3T-FID, Geneva3T-FID, Lausanne3T-ECCENTRIC, Geneva7T-ECCENTRIC, and Vienna7T-FID), each built from that dataset’s own regional profiles via Eq. 7 using the same chimeraLFMIHISIF scale-3 parcellation and Spearman connectivity construction throughout. The five matrices illustrate that this construction generalizes cleanly across five independent sites, sequences, and field strengths without any dataset-specific tuning, each preserving a qualitatively similar block-diagonal and modular organization despite the underlying acquisitions differing in essentially every technical respect.

**Figure 6:**
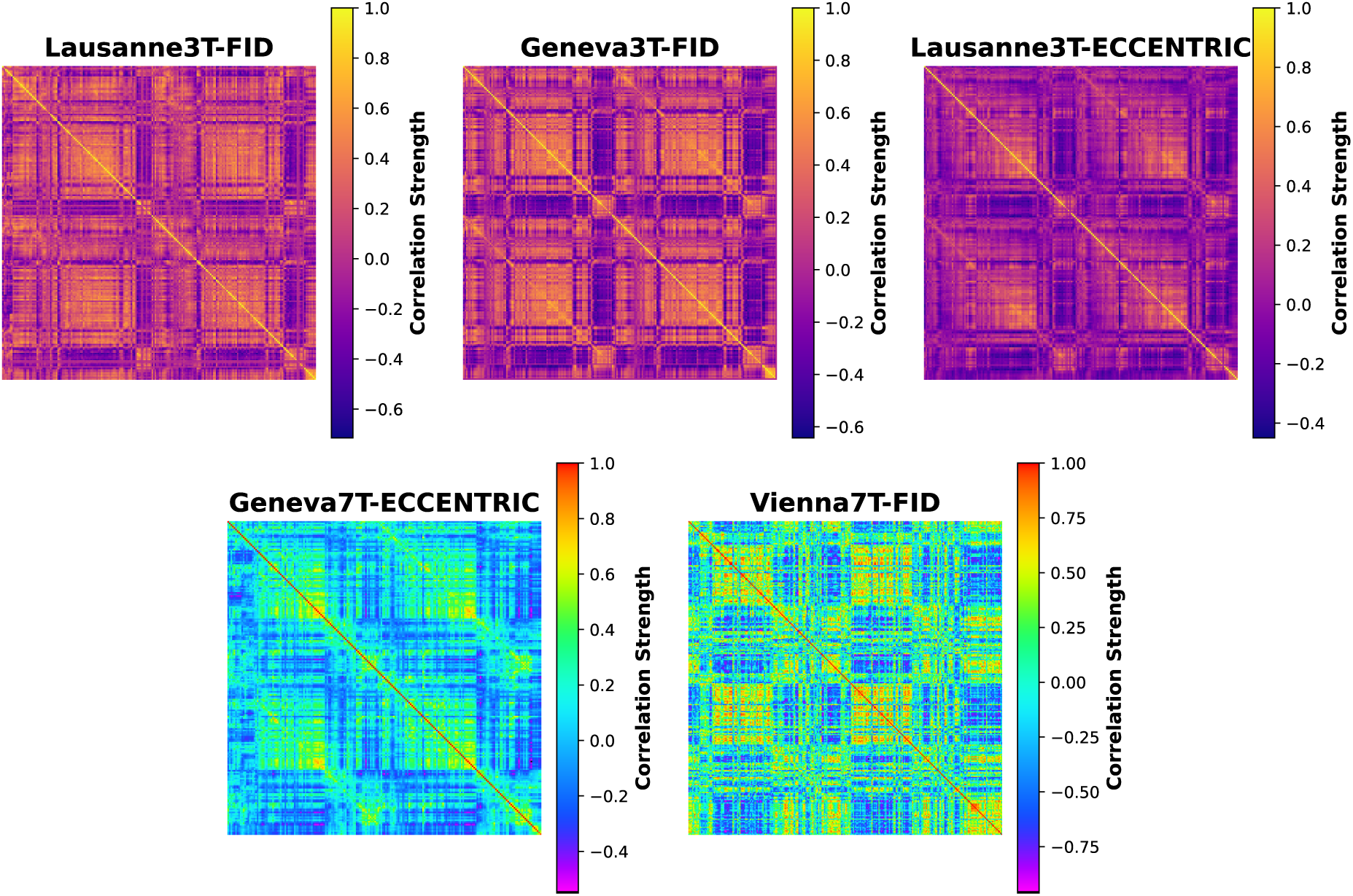
Group-average metabolic connectivity matrices for all five real-world datasets of §3.2. One matrix per acquisition (Lausanne3T-FID, Geneva3T-FID, Lausanne3T-ECCENTRIC, Geneva7T-ECCENTRIC, Vienna7T-FID), constructed identically via Eq. 7. The diagonal is unity by construction; each panel’s color scale (correlation strength) is fit to that dataset’s own dynamic range rather than shared across panels.

## 4 Discussion

We have developed MRSIPrep as a standardized post-quantification workflow for whole-brain MRSI, designed to provide reproducible, modular and transparent processing independently of the downstream analysis. Following software-engineering practices established within the NiPreps ecosystem, MRSIPrep relies on a fixed-version containerized environment, independently executable processing nodes and automated quality-control reports. This architecture enables processing decisions to be explicitly defined and consistently reproduced across datasets and computational environments, while retaining the flexibility required to accommodate different acquisition protocols and analysis strategies.

We evaluated four registration configurations, including ANTs Rigid+SyN, ANTs Rigid+Affine, FSL FLIRT-only and FSL FLIRT+FNIRT, using complementary measures of signal-weighted leakage, detection of a known planted abnormality and gray-matter boundary tracking. We observed that ANTs Rigid+SyN consistently showed the strongest or among the strongest performance across these benchmarks, supporting its implementation as the default registration strategy in MRSIPrep. Furthermore, the regional metabolic profiles obtained after processing showed consistent spatial distributions across three independent sites, five acquisitions, two magnetic field strengths and three MRSI sequences. These findings indicate that the spatial organization recovered by MRSIPrep is preserved across substantially different acquisition settings and is not restricted to a specific scanner, site or sequence.

We further observed that agreement between Lausanne3T-FID and Geneva3T-FID was lower than between Lausanne3T-FID and Lausanne3T-ECCENTRIC, despite the former datasets sharing the same MRSI sequence. In our data, maintaining the acquisition site while changing the sequence therefore resulted in greater agreement than maintaining the sequence across different sites. The present comparisons do not allow the respective contributions of scanner hardware, local protocol implementation, reconstruction procedures or cohort characteristics to be disentangled. Nevertheless, these findings indicate that site-related variability may contribute at least as strongly as sequence-related variability to the replicability of regional MRSI measurements.

The cross-dataset comparisons revealed two practical aspects relevant to multi-site MRSI studies. We observed substantial differences in absolute metabolite levels between datasets, despite preservation of their relative spatial distributions across brain regions. This pattern suggests that a considerable component of between-dataset variability may arise from scanner-, sequence-, reconstruction-or quantification-dependent scaling rather than from differences in the underlying regional metabolic organization. Consequently, although MRSIPrep provides a common processing framework, analyses directly pooling regional metabolite measures across datasets will still require an appropriate harmonization strategy.

An important distinction should nevertheless be made between computational reproducibility and replicability across heterogeneous data. The containerized and version-locked architecture of MRSIPrep ensures that identical inputs and configurations generate identical derivatives independently of the host system. However, computational reproducibility alone does not demonstrate that the resulting derivatives preserve meaningful spatial information, as a processing artifact may itself be perfectly reproducible. The agreement of regional metabolic profiles across independent datasets therefore provides complementary evidence that the spatial patterns obtained after MRSIPrep processing remain stable across differences in acquisition and site. In this context, standardizing post-quantification processing reduces one additional source of variability when comparing results across studies and laboratories.

Several limitations of the present work should be acknowledged. Although validation included proton MRSI acquired at both 3 T and 7 T and using different acquisition strategies, the datasets investigated here do not encompass the full diversity of MRSI sequences, reconstruction approaches and quantification methods currently available. Moreover, MRSIPrep operates downstream of metabolite quantification and therefore cannot correct for biases or errors already introduced during spectral reconstruction or fitting. The reproducibility provided by MRSIPrep consequently applies to the processing steps performed after its input boundary, whereas the reliability of the quantified metabolite maps remains dependent on the upstream reconstruction and fitting framework. Finally, the accuracy of MRSIPrep derivatives remains dependent on the quality and spatial fidelity of the metabolite maps, associated quality metrics and anatomical images provided as input.

Importantly, MRSIPrep operates primarily on quantified spatial maps rather than on sequence-specific k-space or spectral information. Its main processing operations are therefore not intrinsically restricted to proton MRSI. This opens the possibility of extending the framework to other spectroscopic imaging approaches, including ^31^P-MRSI and deuterium metabolic imaging. These applications were not tested in the present study, and differences in signal properties, quantification, quality metrics and biological interpretation may require nucleus-specific adaptations. Nevertheless, we see no fundamental constraint in the current architecture that would prevent such extensions. Rather, the modular and openly available source code provides a foundation on which modality-specific components can be implemented and validated through future contributions from the wider MRSI community.

## 5 Code Availability

The MRSIPrep source code is available under an open-source license at: https://github.com/MRSI-Psychosis-UP/MRSIPrep

Documentation, including the full CLI reference (summarized in Supplementary Table 6), usage examples, and the benchmark and validation results underlying §3, is available at: https://mrsiprep.readthedocs.io/en/stable/

MRSIPrep’s test suite is run via pytest with branch coverage on demand after a push and pull request.

**Software versions**: the results reported here were produced with MRSIPrep v1.9.1, distributed as a Docker image pinning ANTs 2.6.5 [20], FSL 6.0.7.22 [21, 22], FreeSurfer 8.2.0 [23], PETPVC [24], Chimera 0.3.1 [25], and Nipype 1.11.0 [26].

## 6 Data Availability

**Synthetic datasets** (no partcipant data; freely redistributable): the CC0-licensed general-purpose pipeline test dataset introduced in §2.6 (32 subjects, real T1w anatomicals from two CC0 OpenNeuro datasets paired with model-synthesized MRSI signal, no group-level injection) is published as **SynthMRSI-Project** at https://doi.org/10.5281/zenodo.21477047. The synthetic ground-truth cohort underlying the voxel-based-detection benchmark of §3.3 (raw T1w anatomicals and raw/model-synthesized MRSI signal for all 32 subjects, together with the population-level ground-truth injection masks used to score detection performance; per-subject, per-registration-backend processed derivatives are not included, since they are fully regeneratable from these raw files with MRSIPrep itself) is published as **SynthMRSI-VBA-Project** at https://doi.org/10.5281/zenodo.21849051.

**Vienna7T-FID**, the public 7 T CRT-FID-MRSI dataset of §3.2, is available from its original authors at https://doi.org/10.5281/zenodo.4382176 [27].

**Lausanne3T-FID, Lausanne3T-ECCENTRIC, Geneva3T-FID, and Geneva7T-ECCENTRIC** (§3.2), together with the single-subject Geneva3T and Geneva7T acquisitions used for the runtime and registration-accuracy benchmarks (§3.1.1, Supplementary Material), cannot be made publicly available due to participants confidentiality. De-identified derivatives may be shared upon a well-motivated request to the corresponding author, subject to institutional and ethical approval.

## 7 Acknowledgements

This project was supported by the Swiss National Science Foundation (grant number 215728); P.K. was supported by a fellowship from the Adrian *&* Simone Frutiger foundation. The 7T data from Geneva were acquired with the support of the Swiss National Science Foundation (SNSF grants 320030_144260, 320030_179404, and 320030_212476 to S.E.), The NeuroNA foundation, The Copley May foundation and the Synapsy Center – The Synaptic Bases of Mental Diseases. The 3T data from Geneva were acquired with the support of the Leenaards Foundation. The 3T data from Lausanne were acquired with the support of SNSF and the NeuroNA foundation. The study also benefited from the imaging infrastructure and support of the Center for Biomedical Imaging (CIBM), a Swiss research center of excellence founded and supported by CHUV, UNIL, EPFL, UNIGE, and HUG, as well as from the support of the Fondation Pôle Autisme, Geneva. Moreover, we gratefully acknowledge Arnaud Merglen and Camille Marie Piguet for their contribution to the acquisition of the Geneva3T MRSI data.

## 8 Author Contributions

F.L. and E.C. conceptualized the framework, developed the methodology, performed the data analysis, conducted the experiments, contributed to data interpretation, and drafted the initial manuscript. P.H. and A.K. provided critical revisions. R.J. conceptualized the creating of the synthetic MRSI datasets and provided critical revisions. Y.A.-G. performed part of the data preprocessing and provided critical revisions. J.L. contributed to MRI data acquisition and provided critical revisions. S.E. and F.D. provided the Geneva 7T MRI data. P.K. conceptualized the study, supervised the project, and provided critical revisions.

## Supplementary Material

**Table 5:**
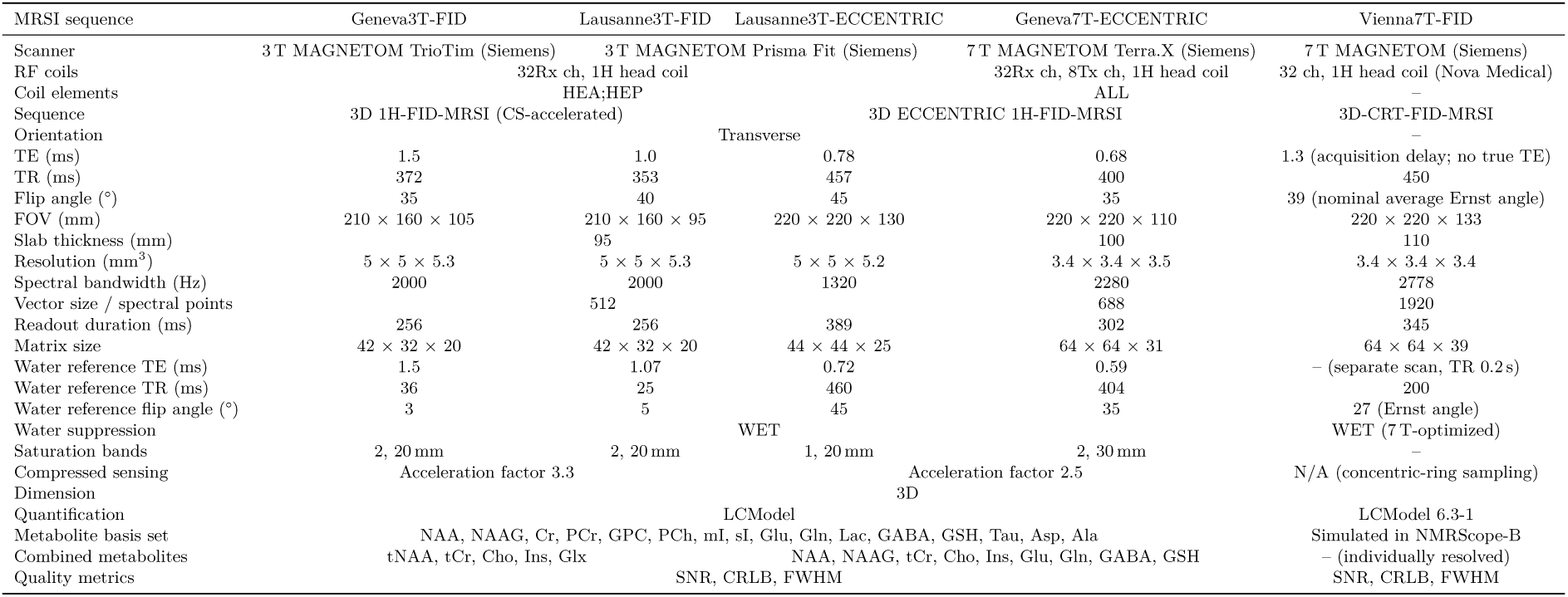
Supplementary Table S1: MRSinMRS acquisition parameters. [17] for the five multi-site datasets underlying the cross-site/cross-sequence regional profile replicability comparisons of §3.2. Geneva3T, Lausanne3T (2019–2022), Lausanne3T (2025–2026), and Geneva7T parameters are drawn from the acquiring sites’ own MRSinMRS reporting; Vienna7T parameters are drawn from its public mrsinmrs.json sidecar, following Hangel et al. [27].

Summary Spearman *ρ*/*p*-values for the cross-site/cross-sequence regional profile comparisons of §3.2 are reported in Table 2; the four figures below show the underlying region-by-region profiles those correlations are computed from, one metabolite per panel, split across four page-sized figures.

**Figure 7:**
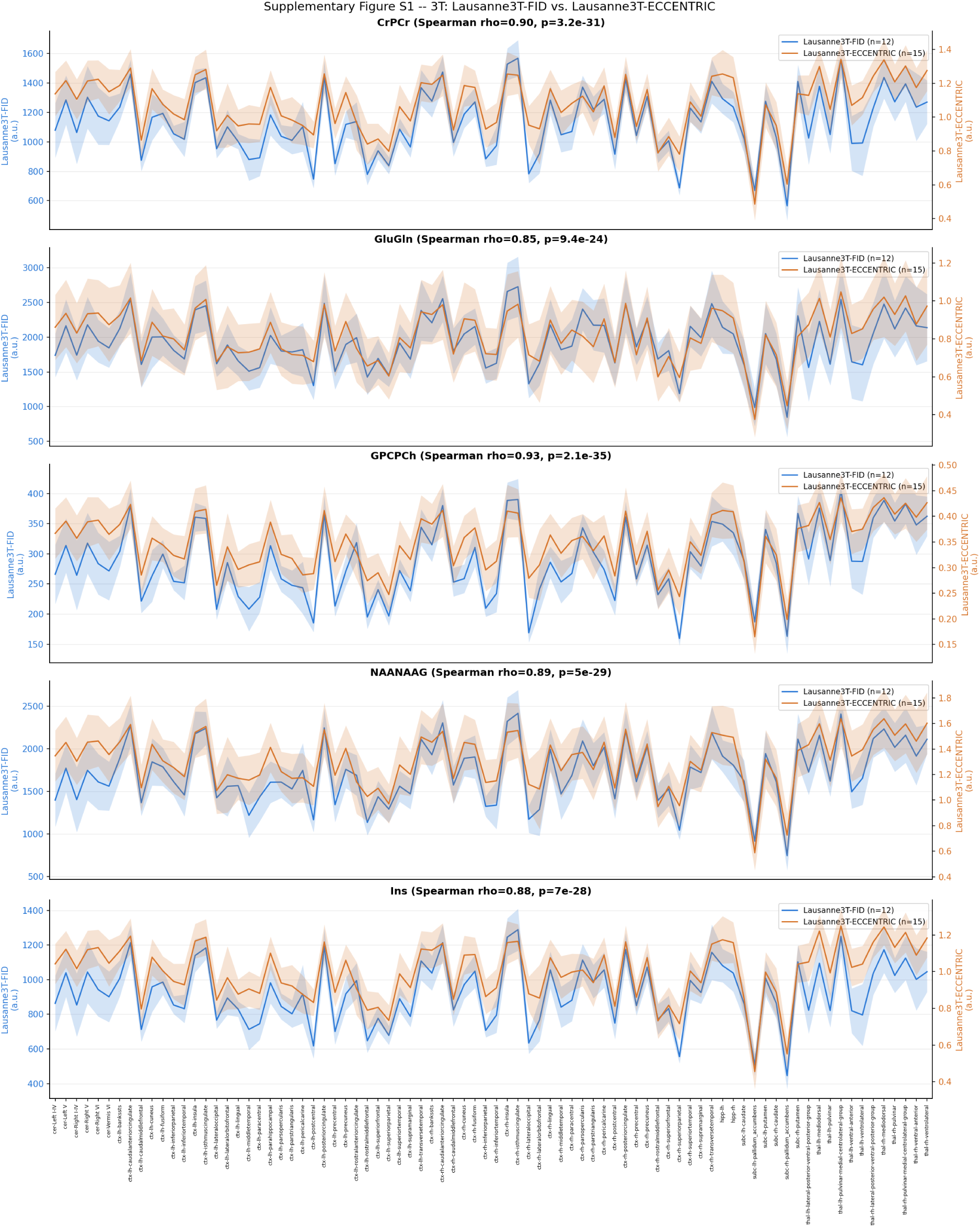
Supplementary Figure S1: 3 T regional metabolic profile replicability, full per-region trace plots. Lausanne3T-FID vs. Lausanne3T-ECCENTRIC, 82 GM root regions, all 5 metabolites. Each dataset is plotted on its own *y*-axis (left: Lausanne3T-FID; right: Lausanne3T-ECCENTRIC) since the two acquisitions’ absolute signal levels differ by roughly three orders of magnitude (a scanner/sequence/ reconstruction scaling difference, not a data error, see §3.2); shaded bands are ±95% CI across subjects within each cohort. Spearman *ρ* in each panel title is computed on the two cohorts’ native scales, before this per-axis rescaling for display.

**Figure 8:**
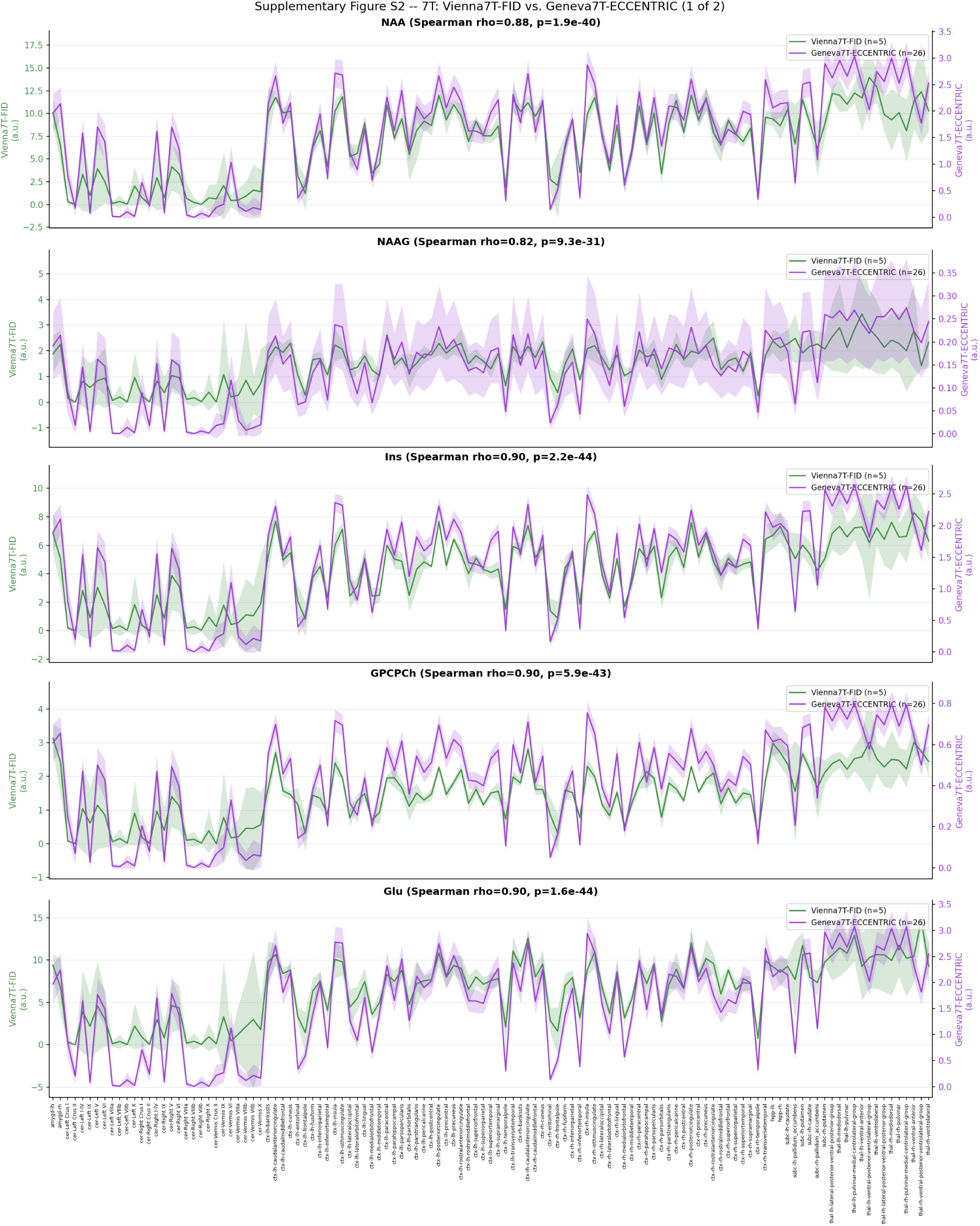
Supplementary Figure S2: 7 T regional metabolic profile replicability, full per-region trace plots (1 of 2). Vienna7T-FID vs. Geneva7T-ECCENTRIC, 119 GM root regions; NAA, NAAG, Ins, GPCPCh, and Glu (remaining metabolites in Supplementary Figure 9). Same twin-axis convention as Supplementary Figure 7 (left: Vienna7T-FID; right: Geneva7T-ECCENTRIC).

**Figure 9:**
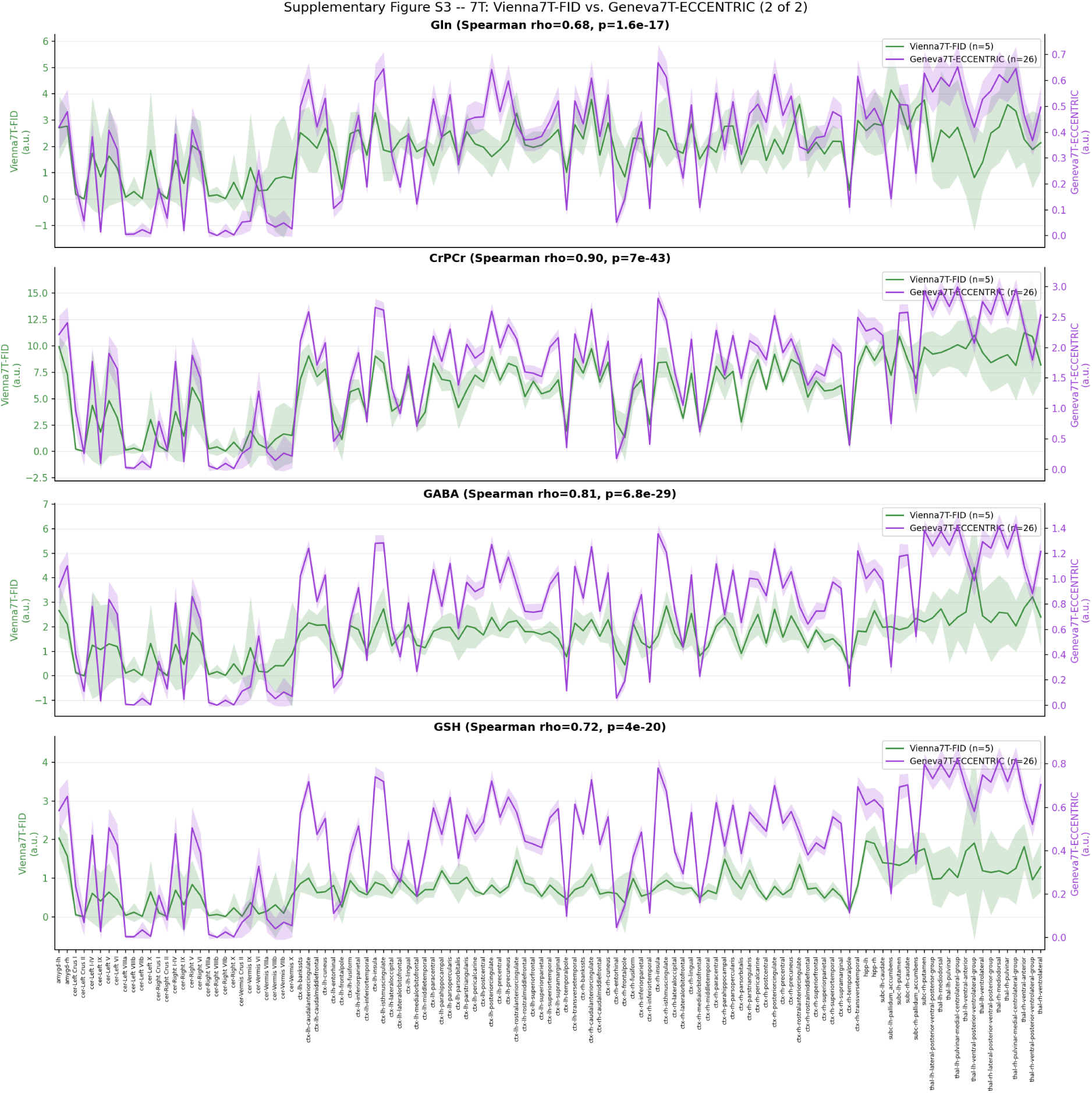
Supplementary Figure S3: 7 T regional metabolic profile replicability, full per-region trace plots (2 of 2). Vienna7T-FID vs. Geneva7T-ECCENTRIC, 119 GM root regions; Gln, CrPCr, GABA, and GSH (remaining metabolites in Supplementary Figure 8). Same twin-axis convention as Supplementary Figure 7.

**Figure 10:**
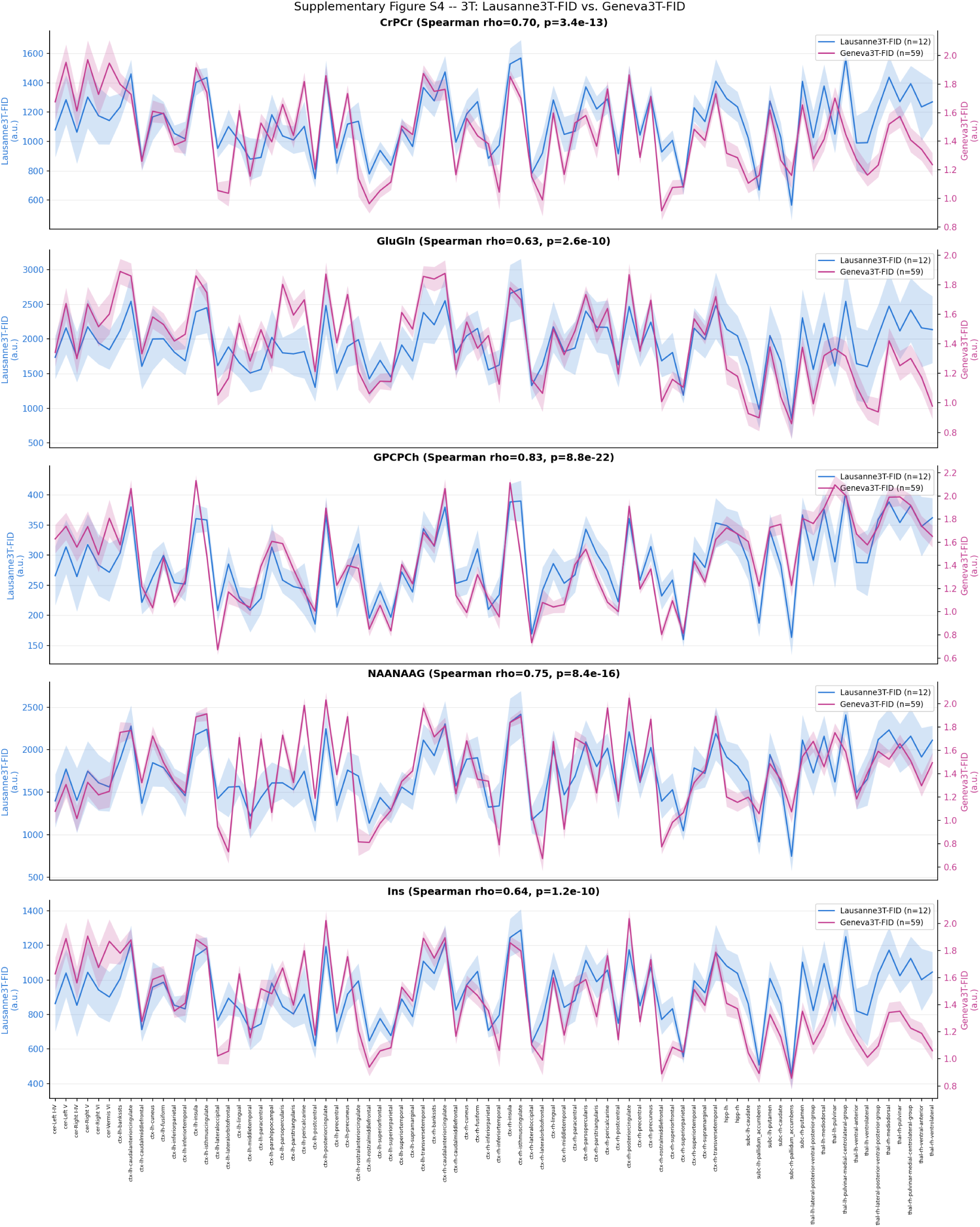
Supplementary Figure S4: 3 T regional metabolic profile replicability, full per-region trace plots. Lausanne3T-FID vs. Geneva3T-FID, 82 GM root regions, all 5 metabolites. Same twin-axis convention as Supplementary Figure 7 (left: Lausanne3T-FID; right: Geneva3T-FID).

### Runtime Scaling

Full –parcellation-mode synthseg runs of both real subjects of §2.6 (Geneva3T and Geneva7T) were repeated at –nthreads 8, 12, 16, and 32 (–nproc 1, one subject per run, executed strictly sequentially with no concurrent load and a fresh –work-dir per run, so every timing reflects a genuine full-pipeline computation rather than a partially cached rerun). **Figure 11** shows total wall-clock time and its breakdown by pipeline step.

**Figure 11:**
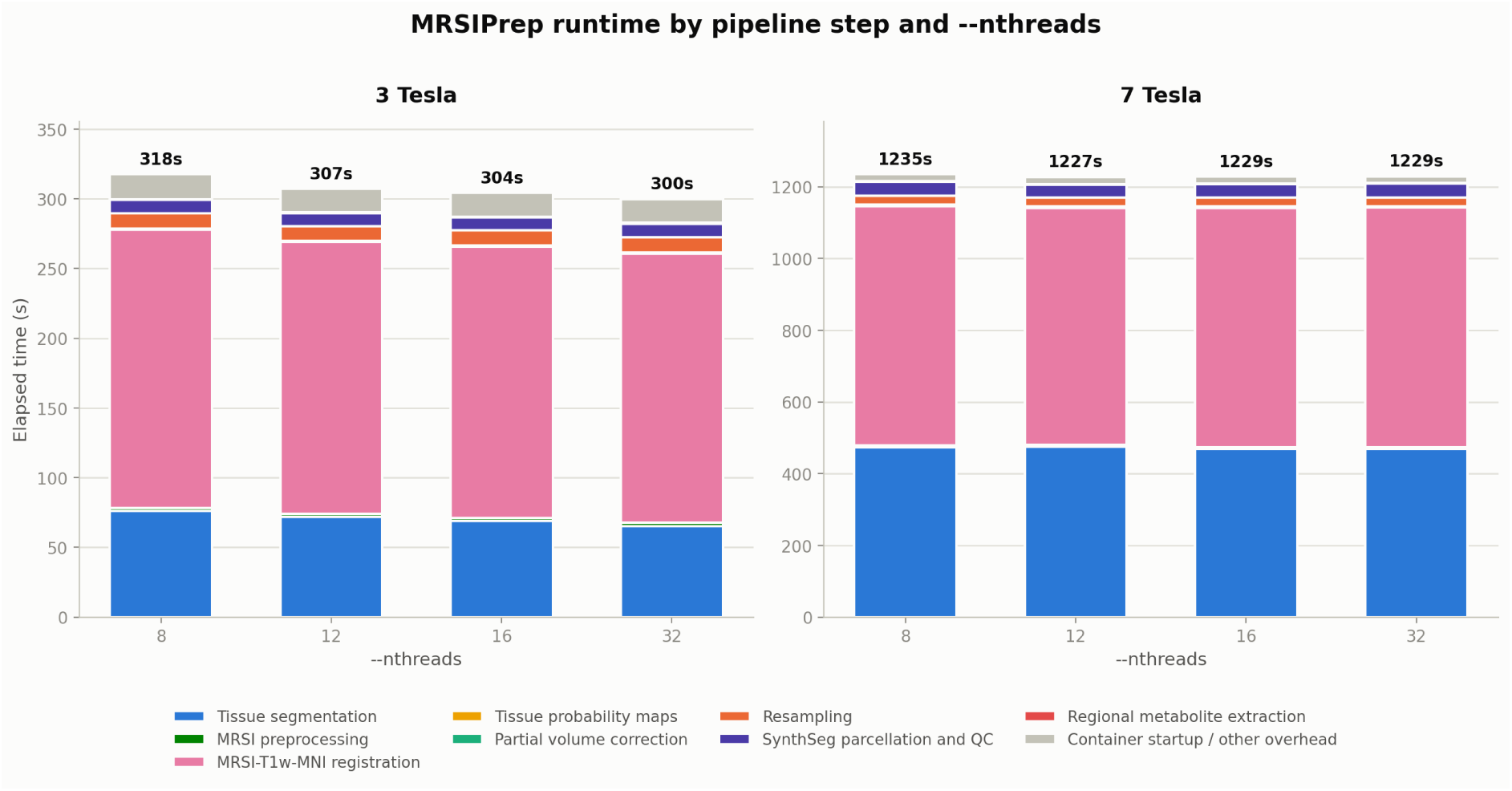
Runtime by pipeline step and –nthreads, 3 T vs. 7. **T.** Stacked bar height is total wall-clock elapsed time (labeled above each bar); segments show each step’s share. “Container startup / other overhead” covers Docker startup and the CLI’s own preflight input check.

Runtime is essentially flat from 8 to 32 threads at both field strengths (3 T: 300–318 s, ∼6% spread; 7 T: 1227–1235 s, under 1% spread), since ANTs registration and SynthSeg tissue segmentation (together the majority of total runtime) show no consistent benefit from more than ∼8 threads; for batch processing, allocating ∼8 threads per subject and increasing –nproc is therefore preferable to allocating more threads to a single subject. The 7 T subject takes ∼4× longer than the 3 T subject (∼20.5 vs. ∼5.1 min), driven primarily by its ∼7.2× larger T1-weighted volume (256×396×416 vs. 160×192×192 voxels) rather than by MRSI grid size, which differs by only ∼2.1× (32,638 vs. 15,315 useful voxels). Anatomical, not MRSI, resolution is the dominant runtime driver, since ANTs registration and SynthSeg both operate on the full-resolution T1w volume.

### Command-Line Interface Reference

MRSIPrep exposes 3 required positional arguments and 78 optional flags across 10 functional groups (Table 6); only –metabolites and –ref-met are required beyond the positionals. The 8 per-step –overwrite-* flags (identical in form and purpose, one per cacheable pipeline stage) are condensed into a single row. The full, auto-generated reference — one entry per flag, always in sync with the argument parser — is published at https://mrsiprep.readthedocs.io/en/stable/usage_basic.html.

**Table 6:**
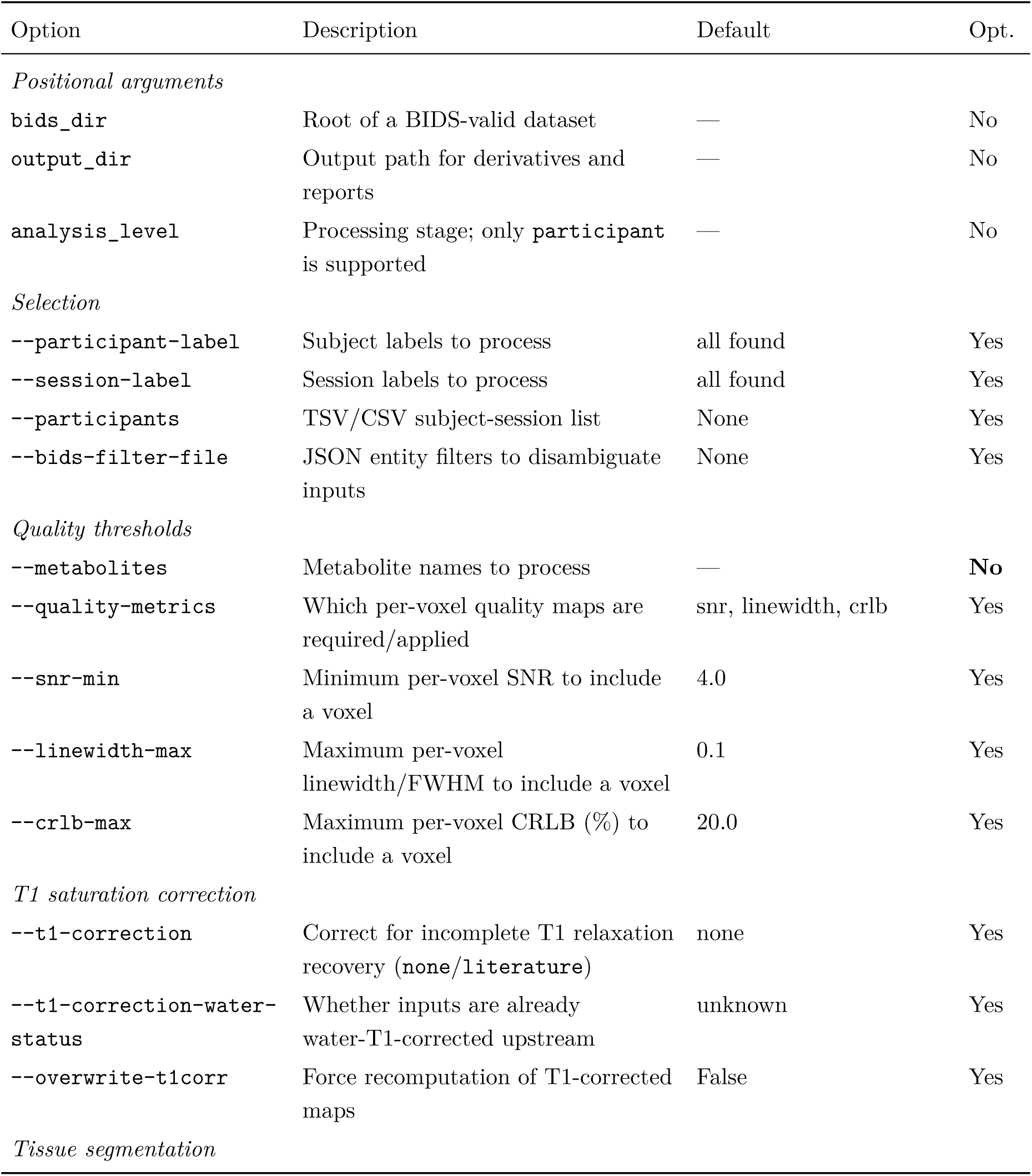
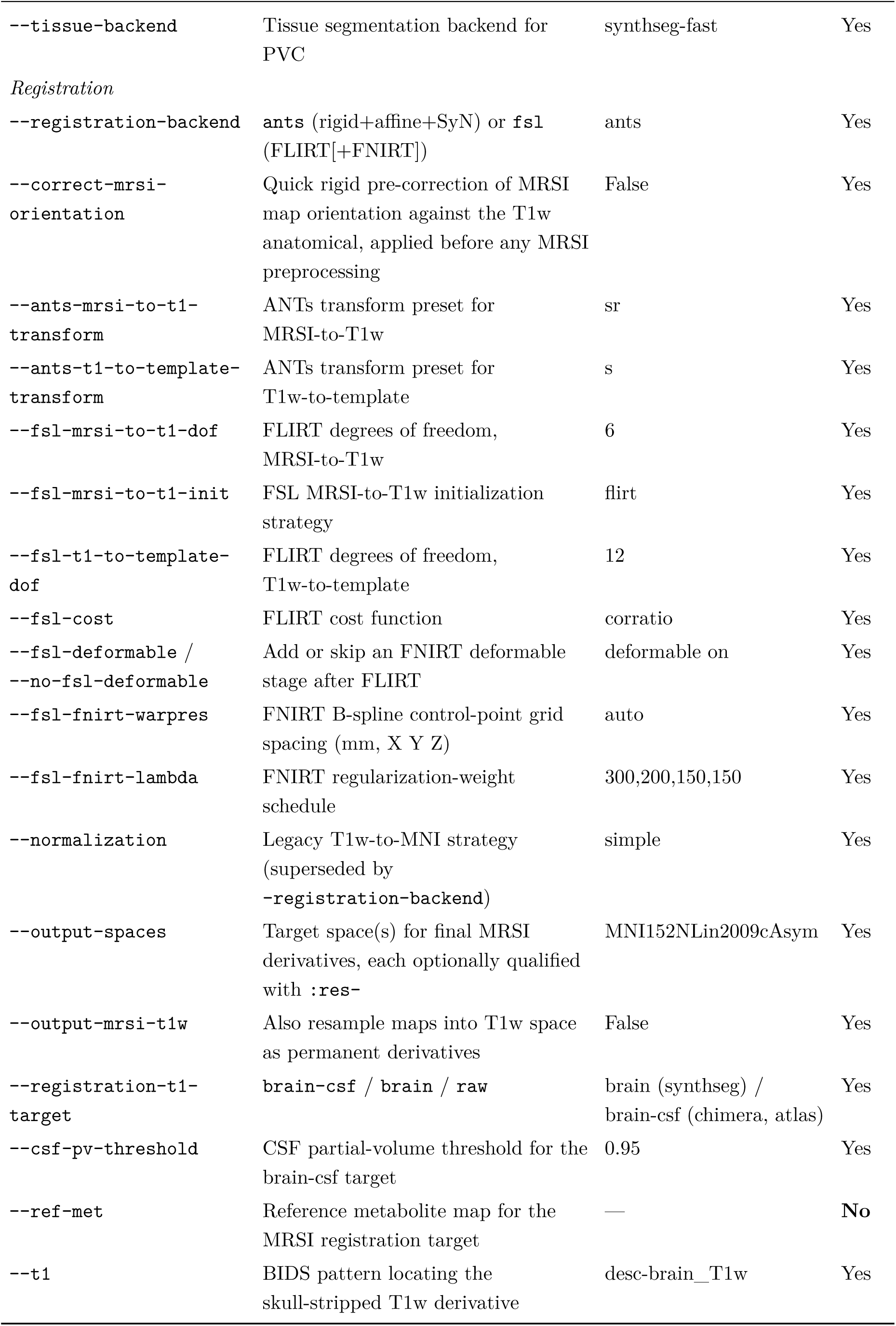
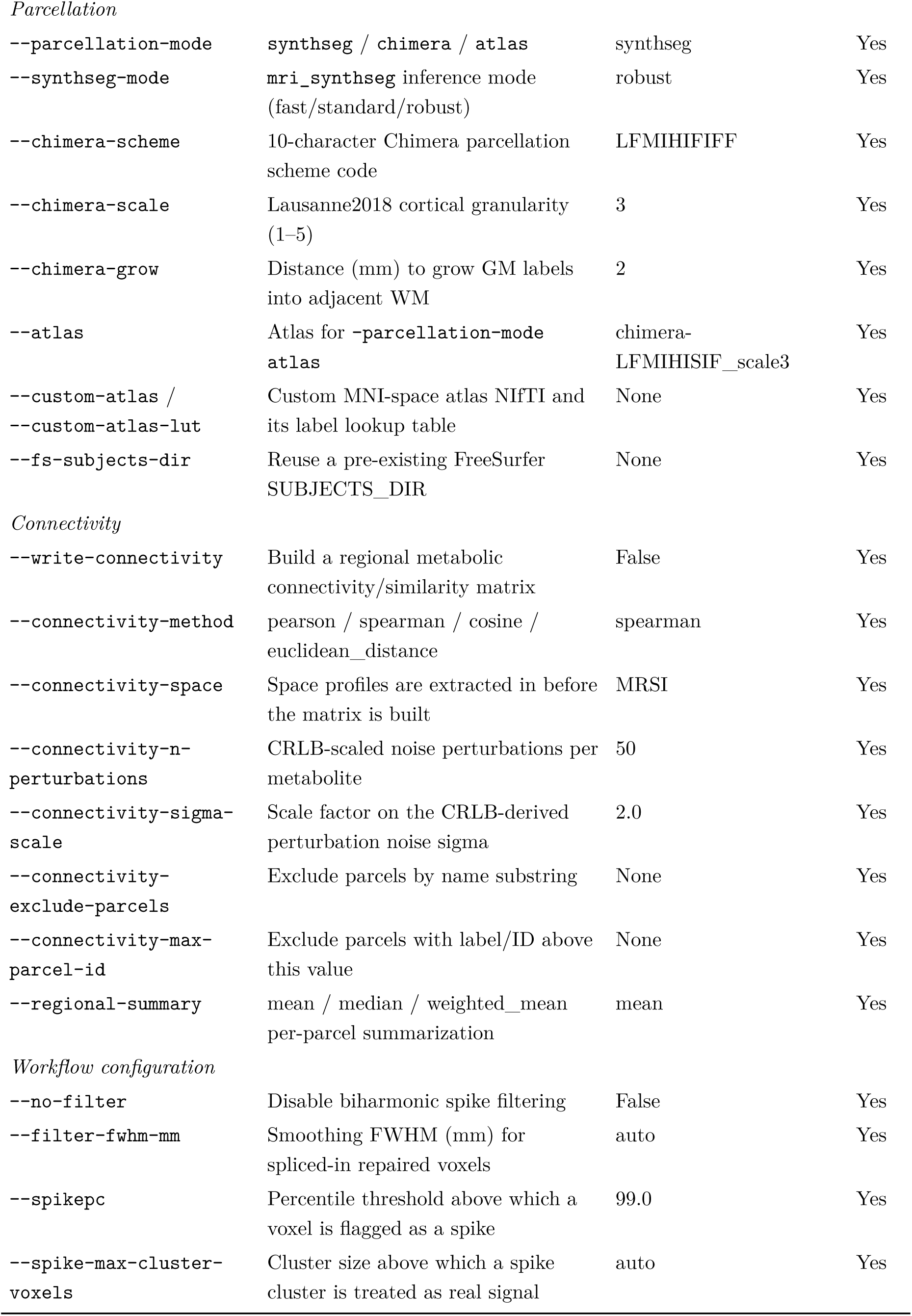
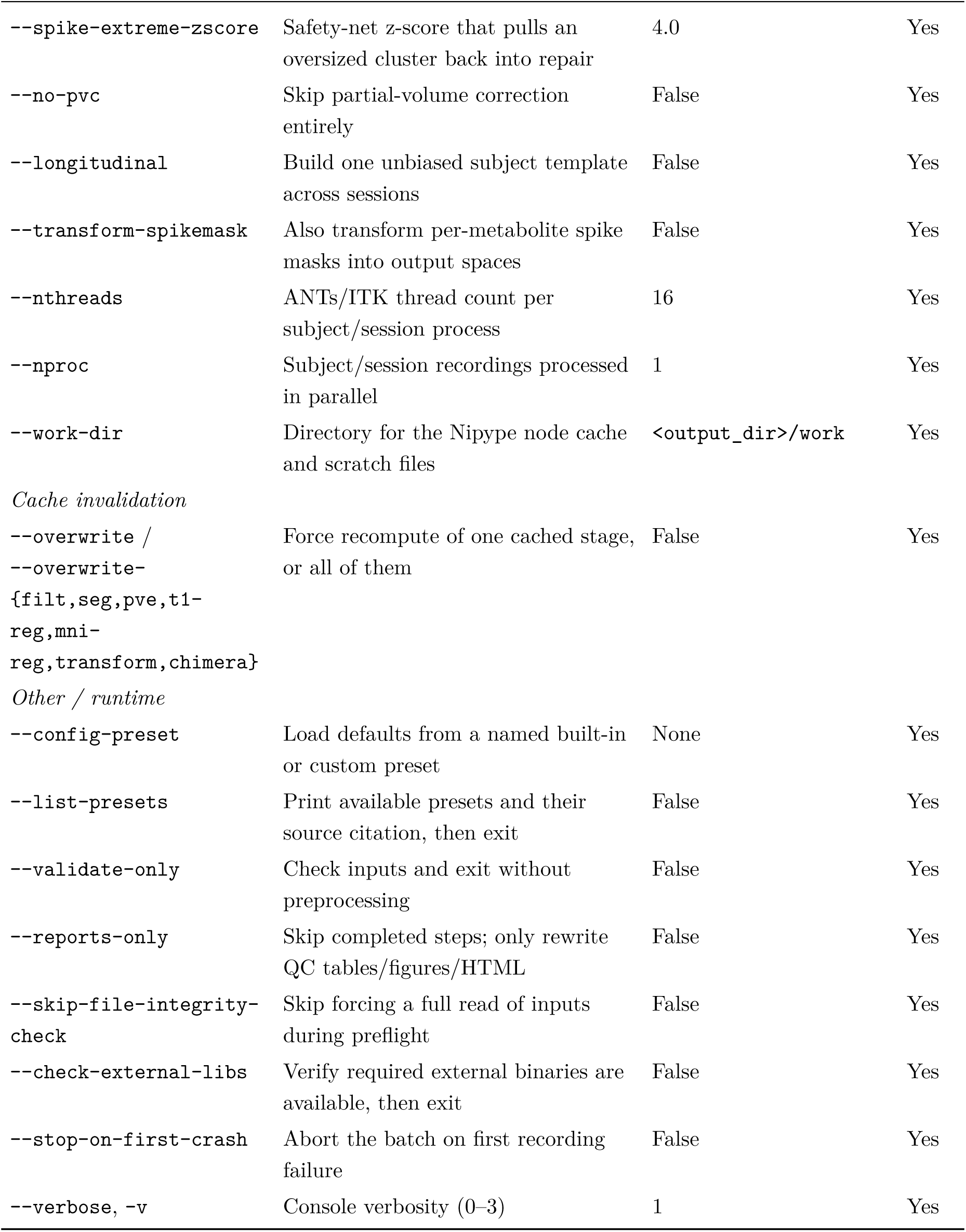
Supplementary Table S2: MRSIPrep command-line options. Option, one-line description, default value, and whether the flag is optional.

### Supplementary Report Example

The following pages reproduce, in full, the automated per-recording QC report MRSIPrep generates for a subject from the public SynthMRSI-Project dataset (§2.6), run end-to-end with –parcellation-mode chimera –write-connectivity. Every tab of the interactive HTML report (normally navigated one at a time in a browser) is expanded here as its own section. The live, interactive version – along with its figures and full provenance record – is available at https://github.com/MRSI-Psychosis-UP/MRSIPrep/tree/main/docs/examples/sub-01_ses-01.

#### MRSIPrep report: SynthMRSI-Project · sub-01 ses-01

##### Inputs

BIDS directory: /data

Output directory: /out/mrsiprep

Parcellation mode: chimera

Tissue backend: synthseg-fast

##### MRSI Raw QC

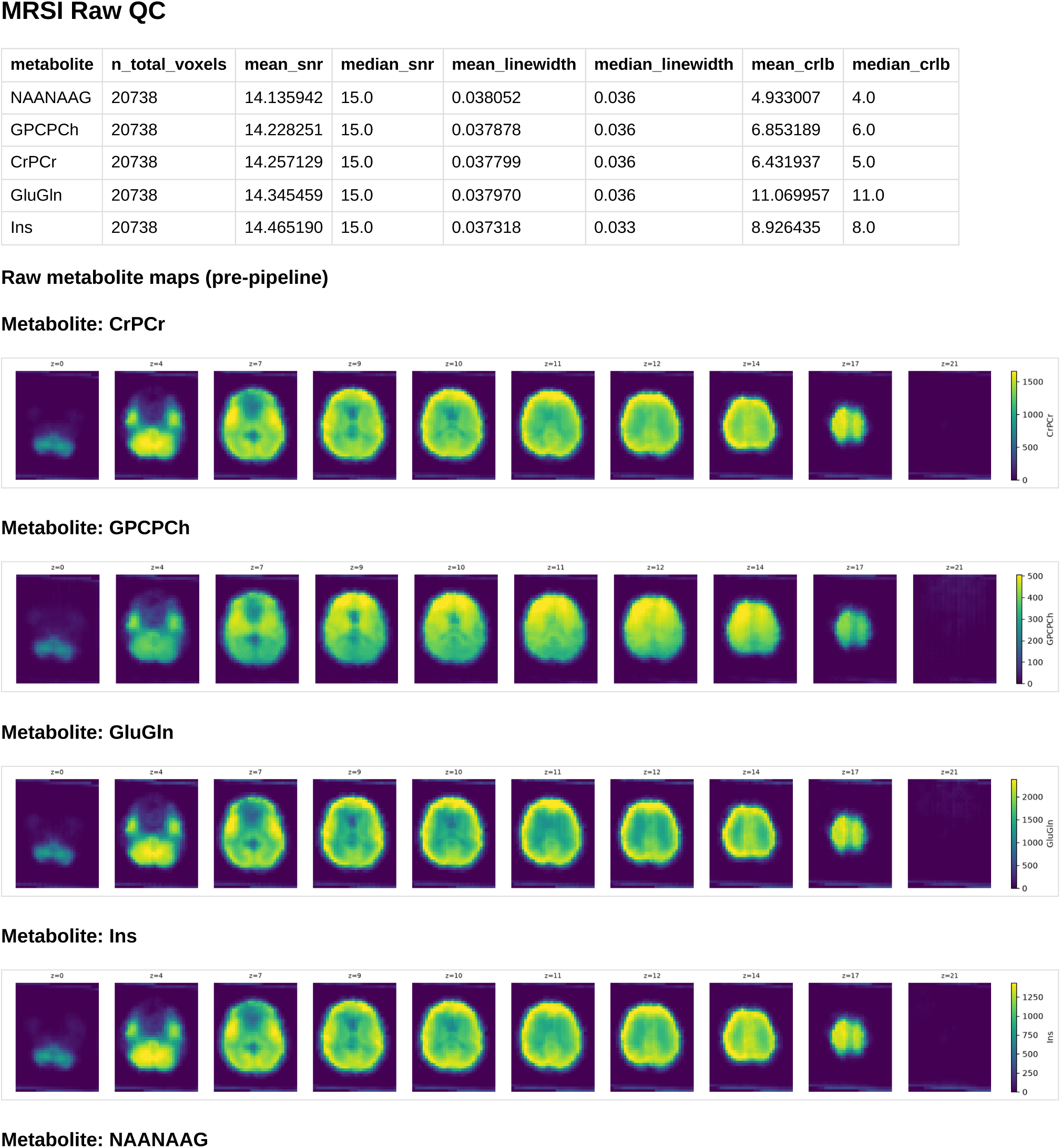
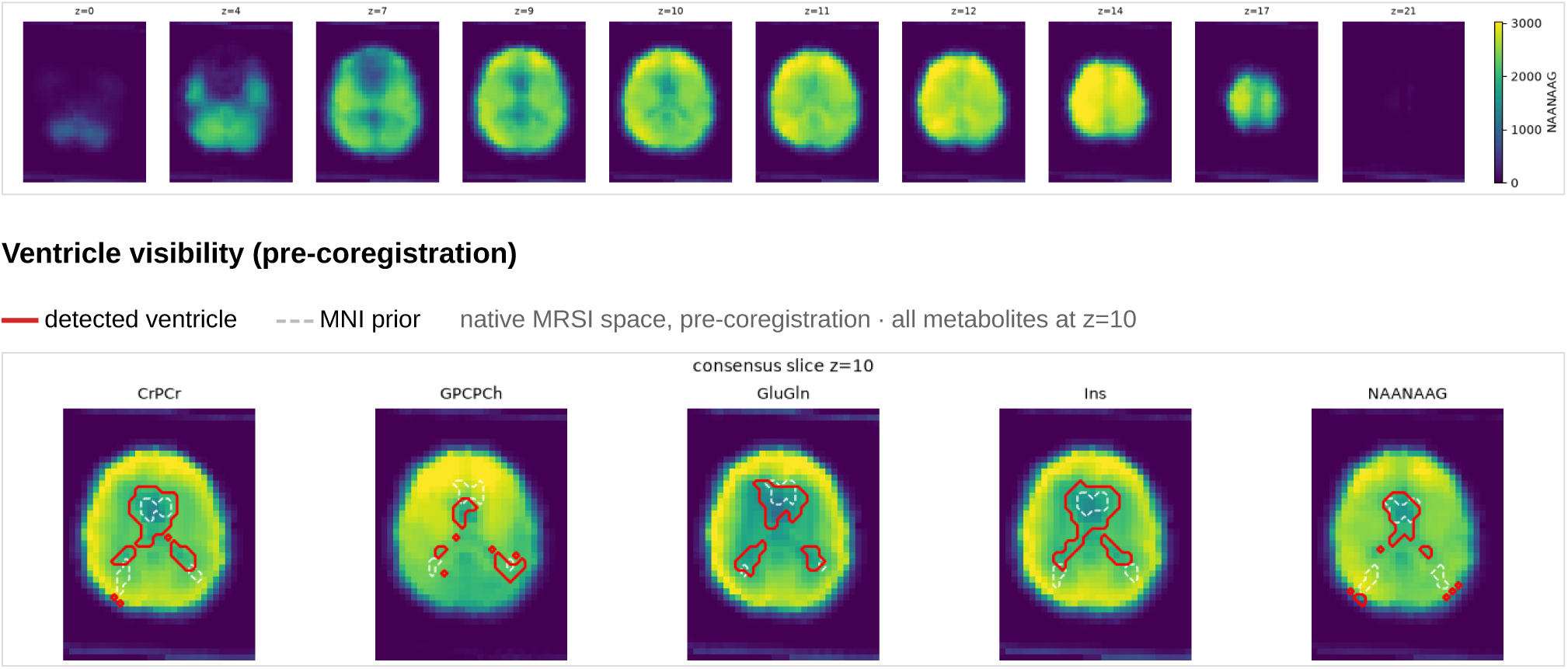

##### MRSI PVC

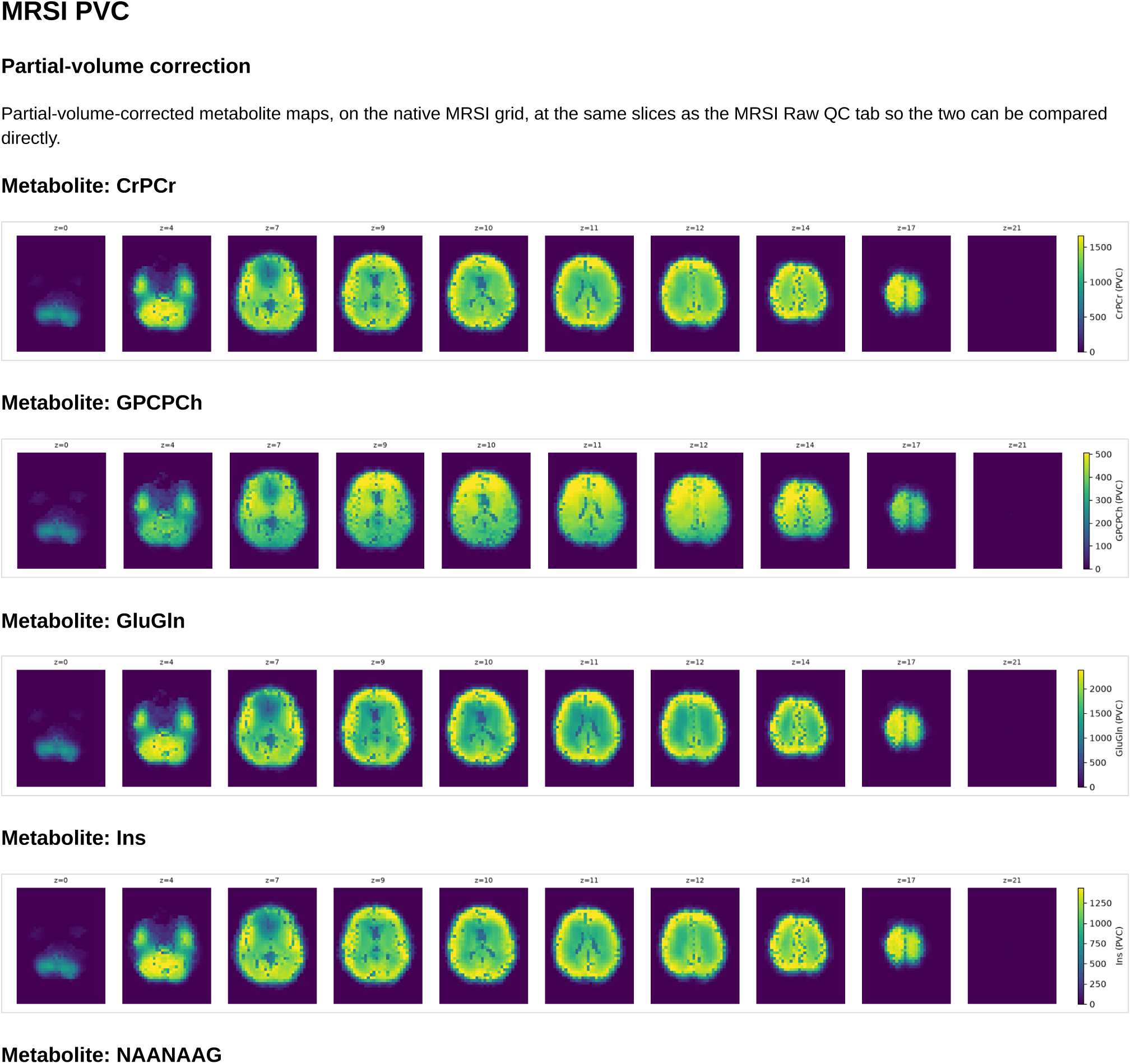
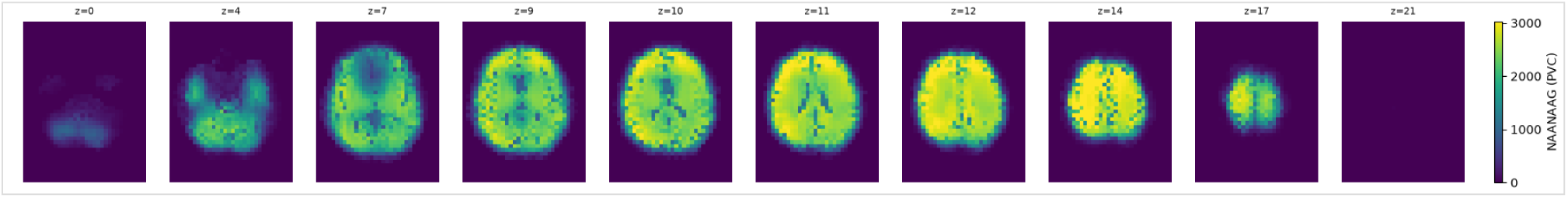

##### Spike Filter

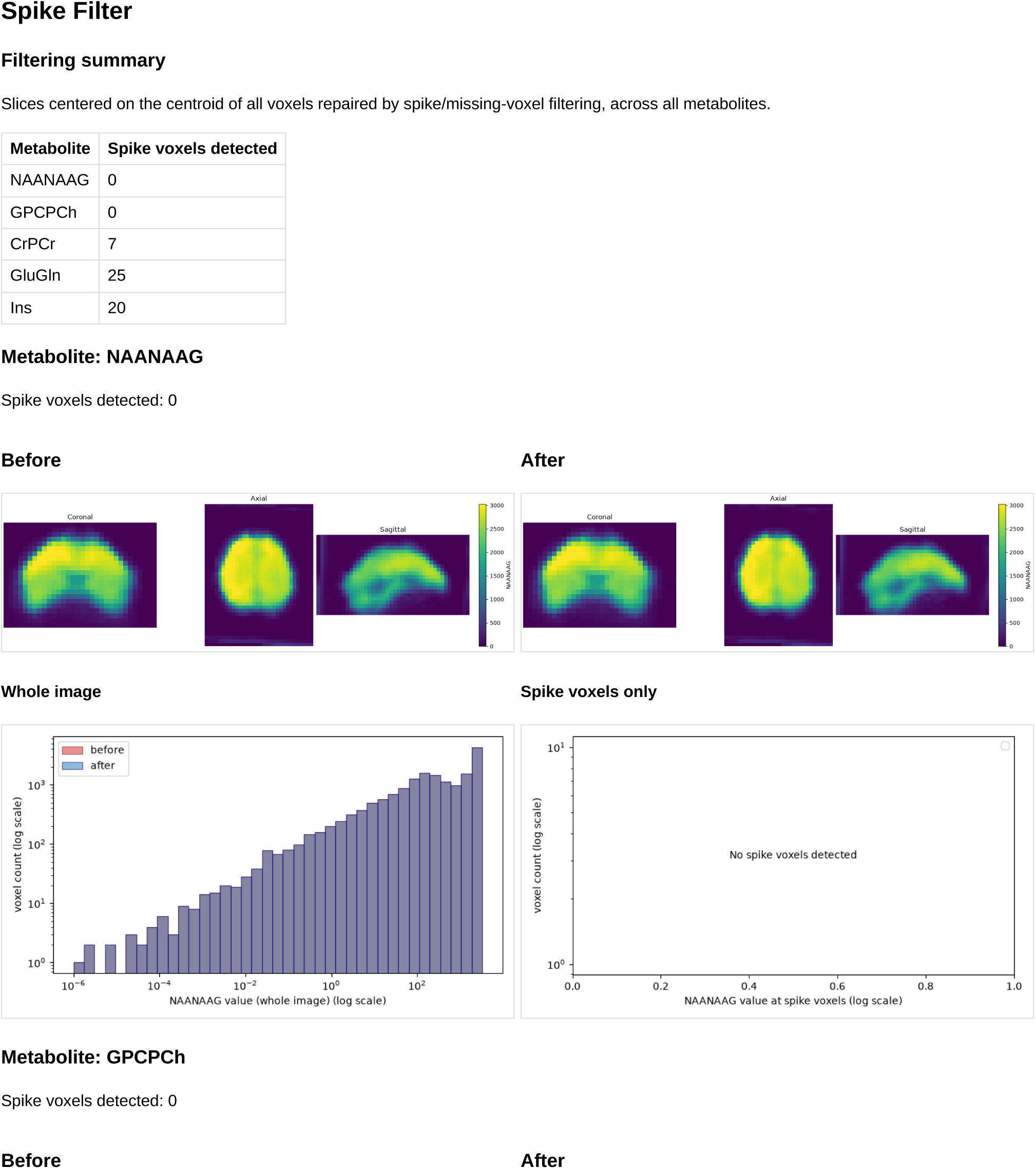
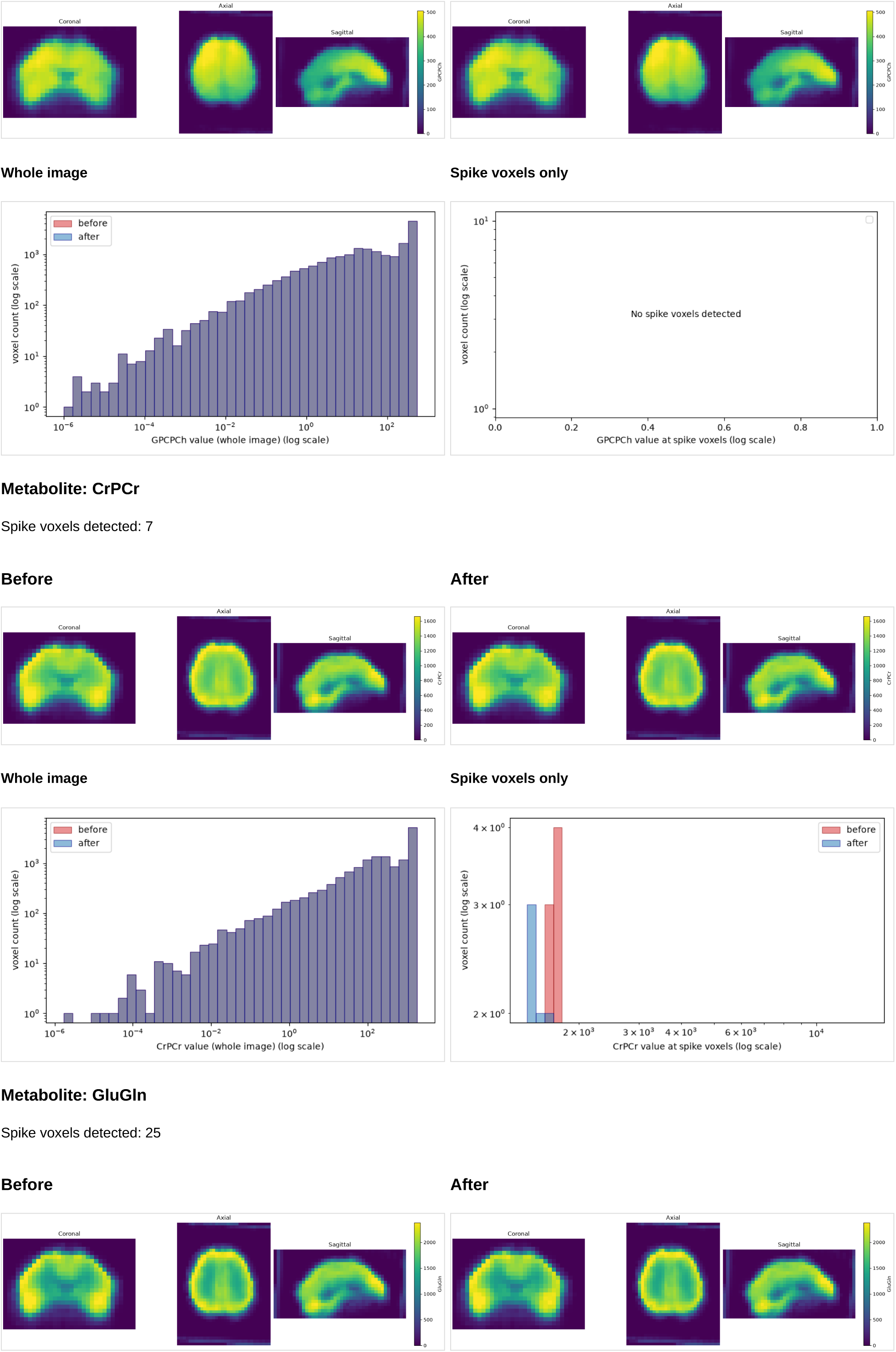
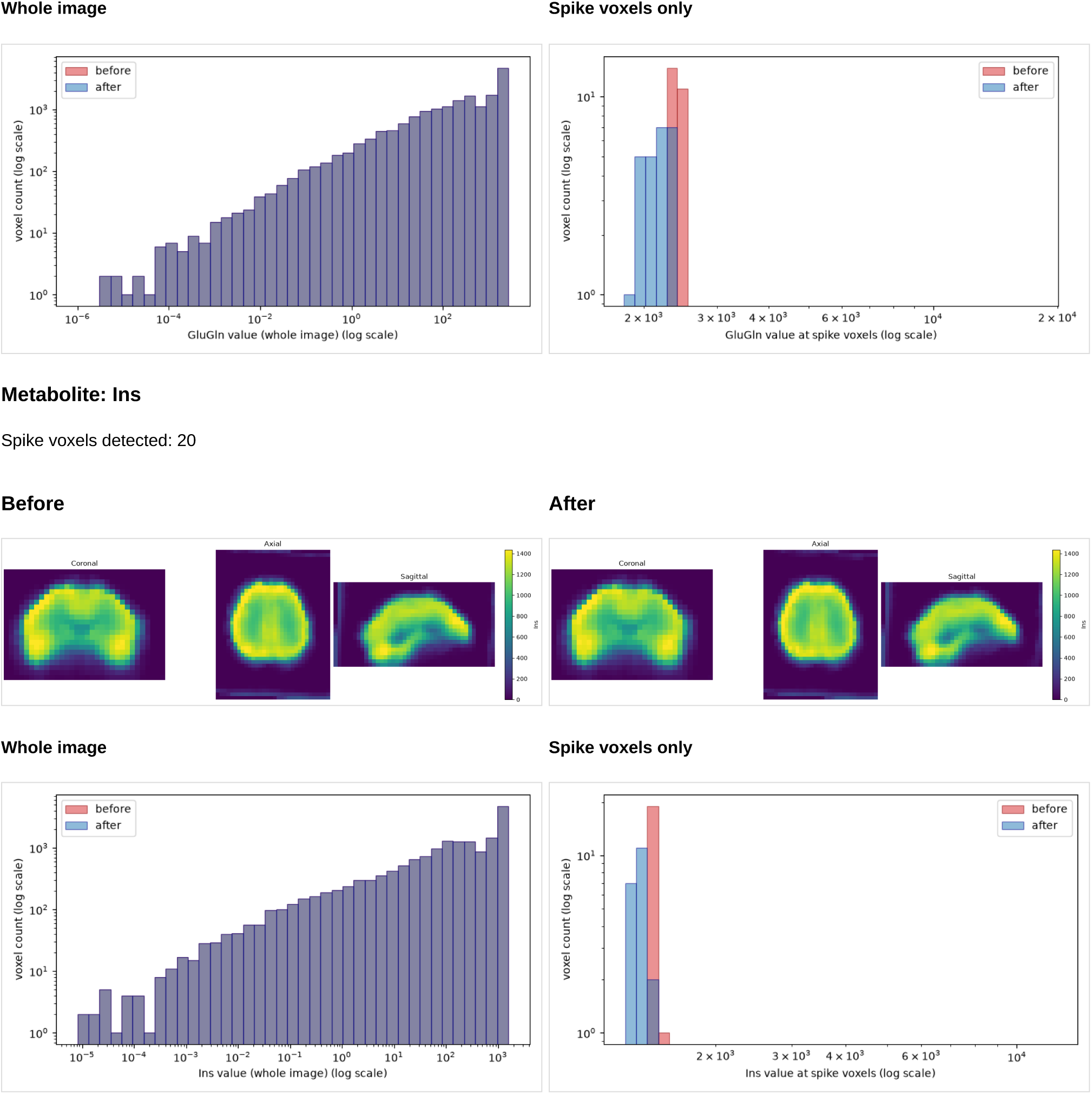

##### Anatomical

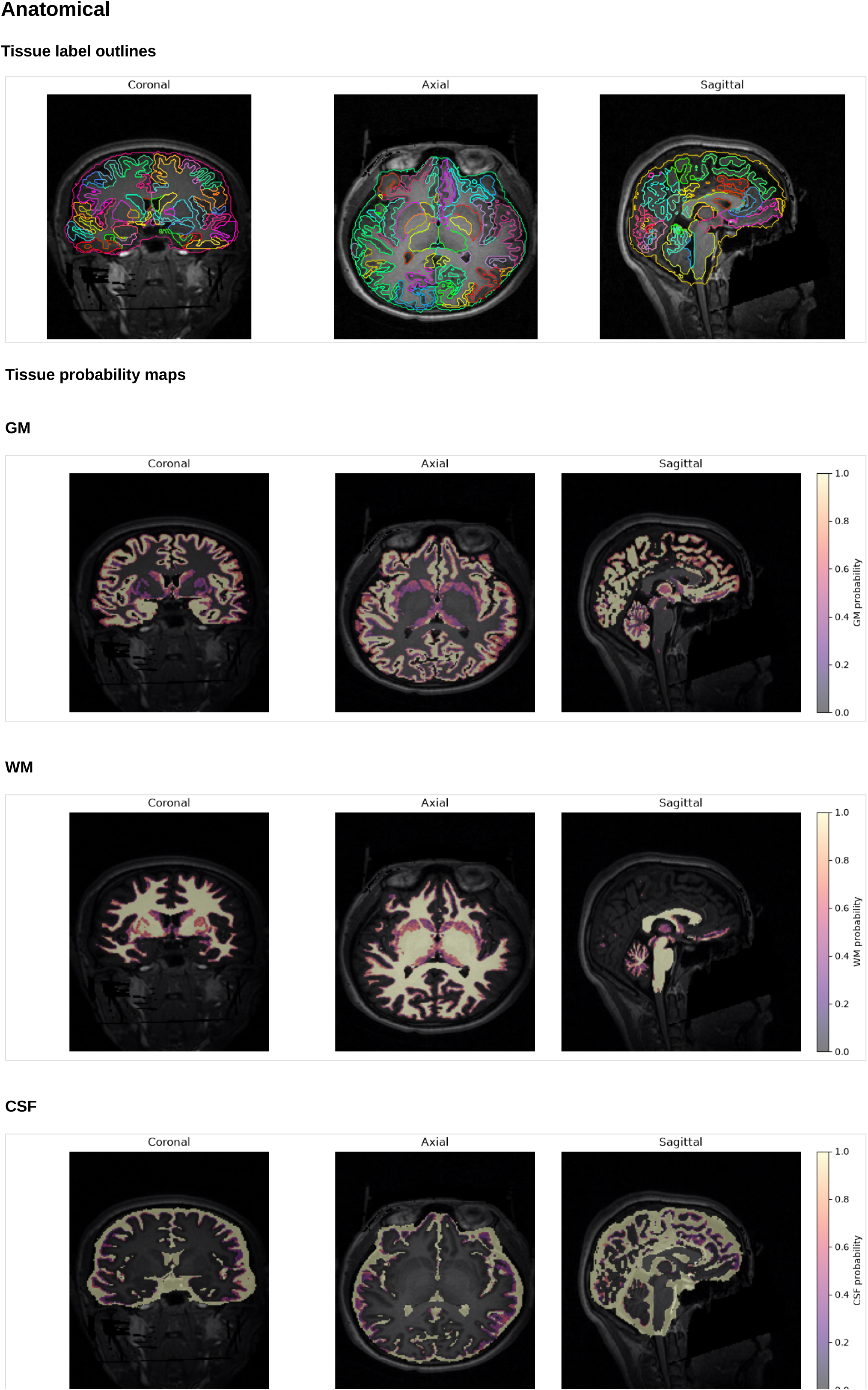

##### T1-space alignment

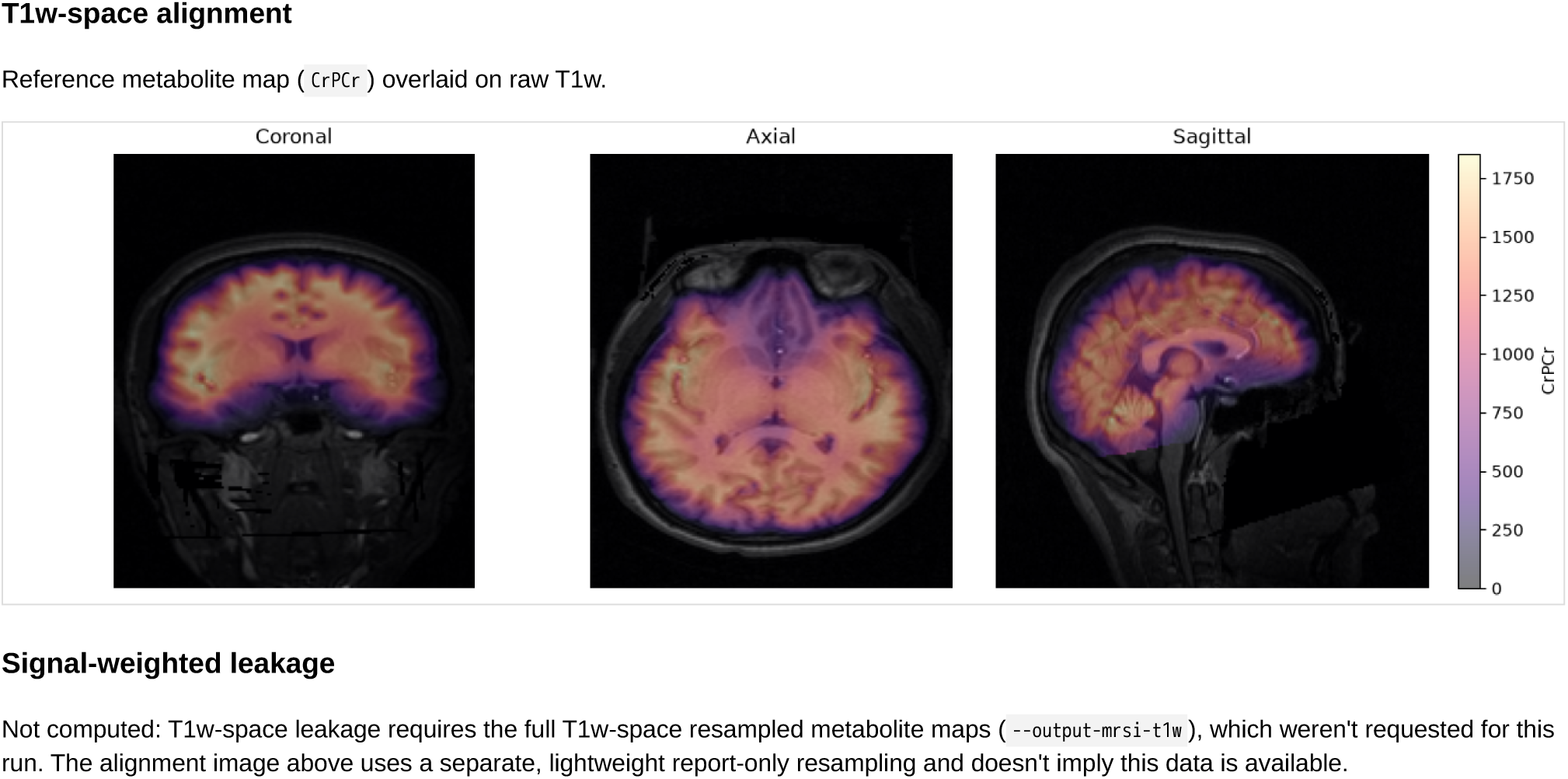

##### Template-space alignment

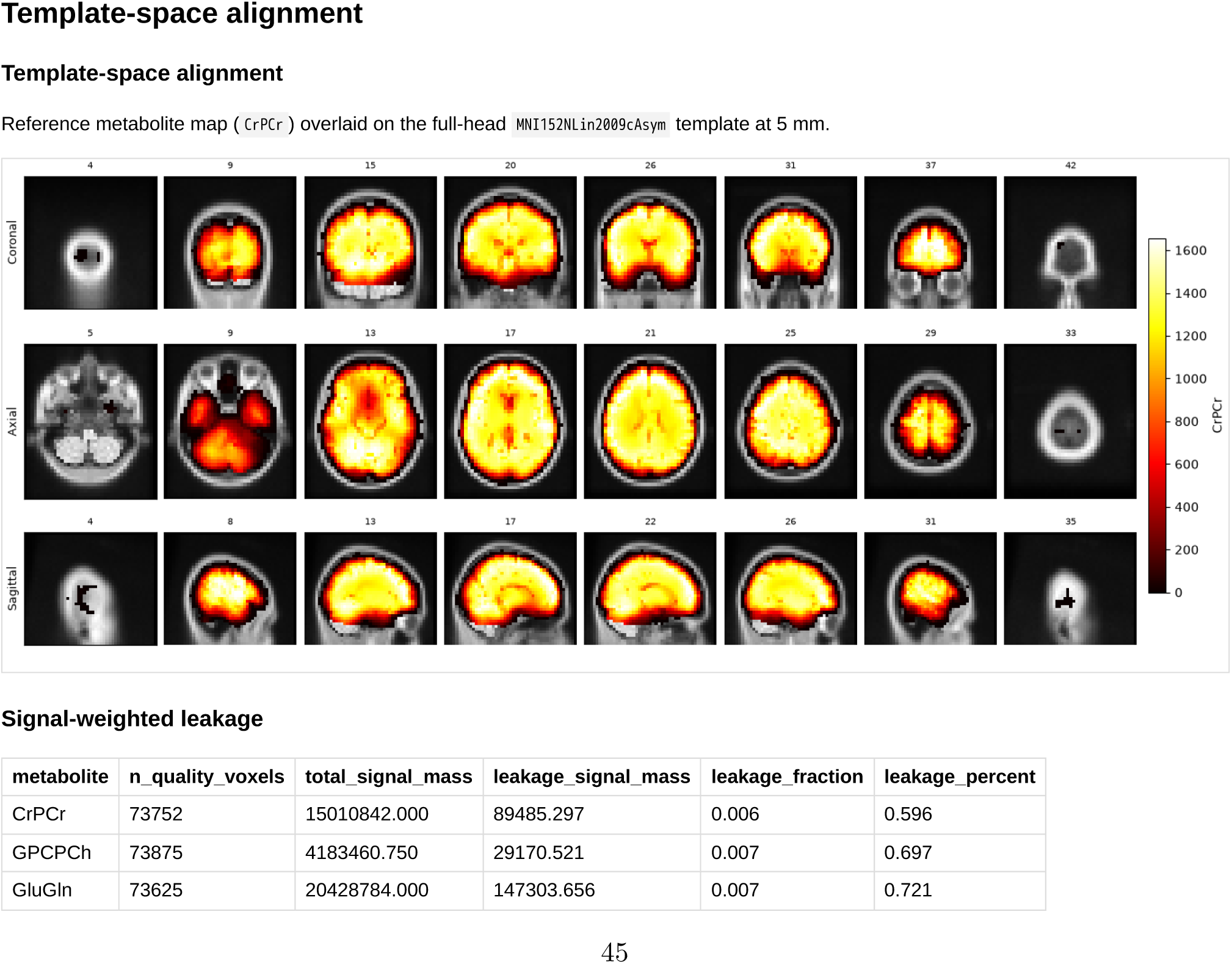
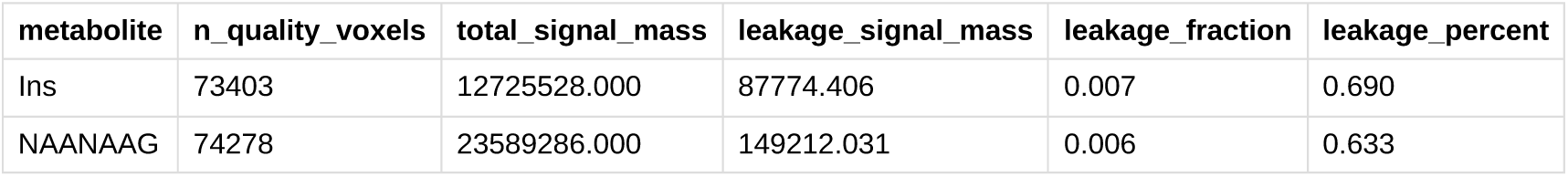

##### Coverage

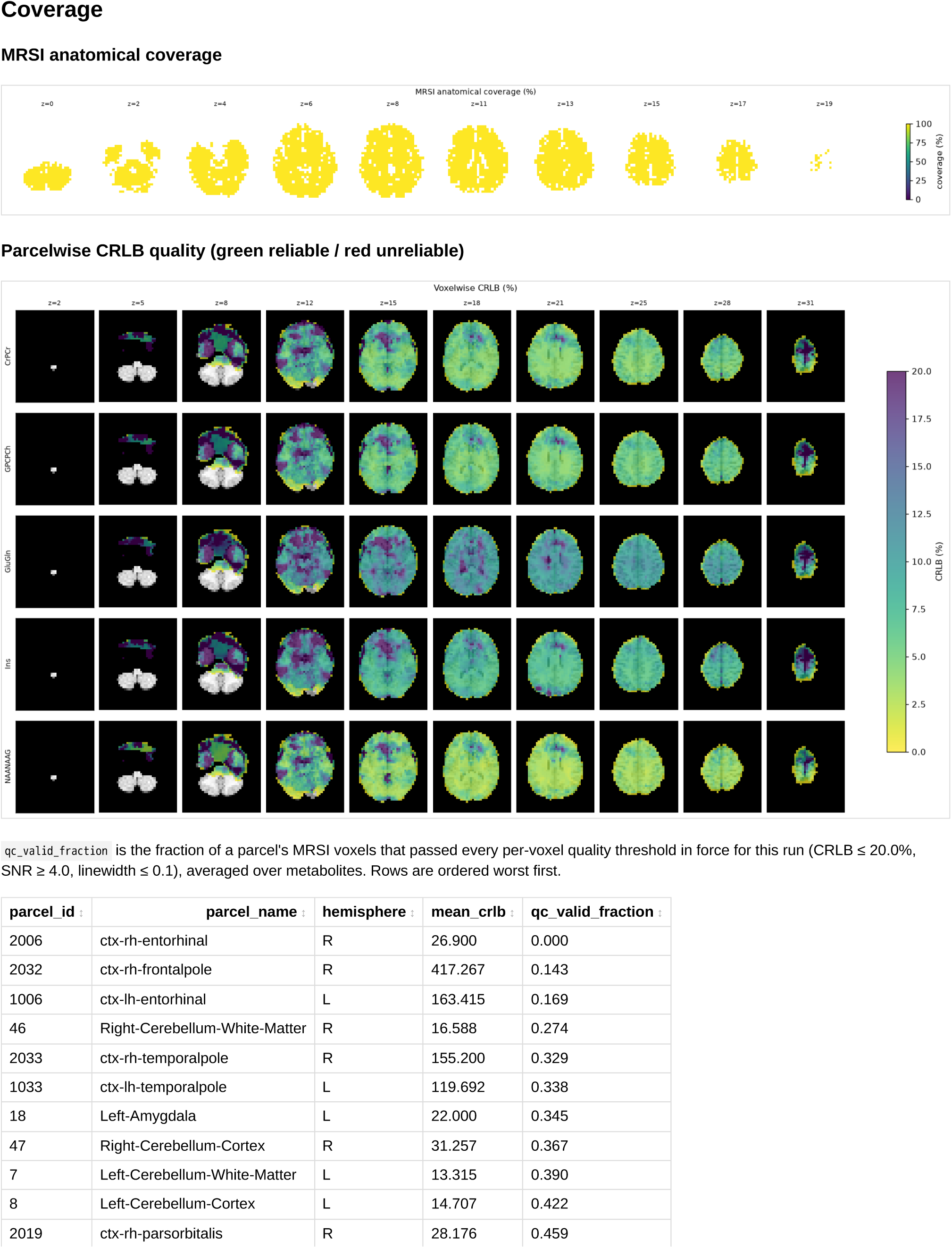
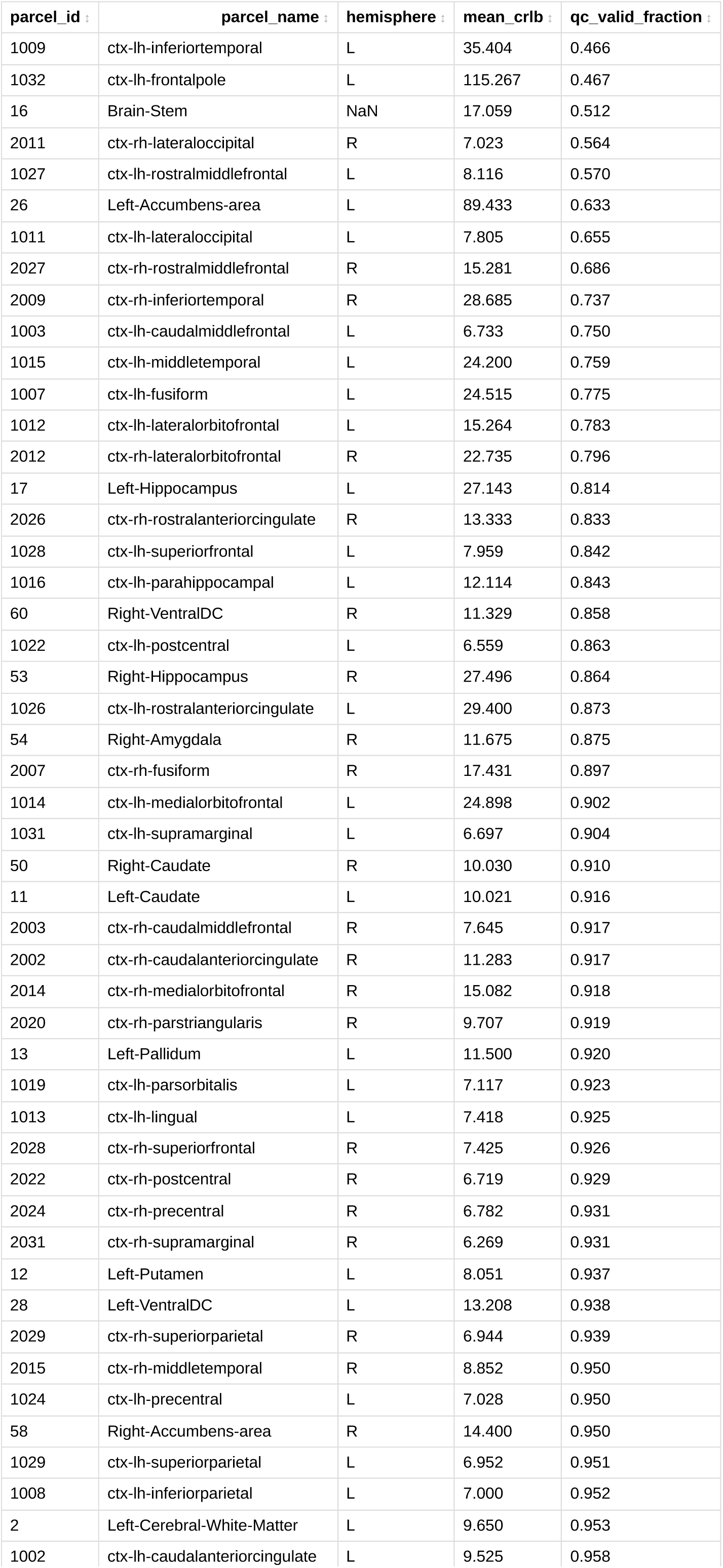
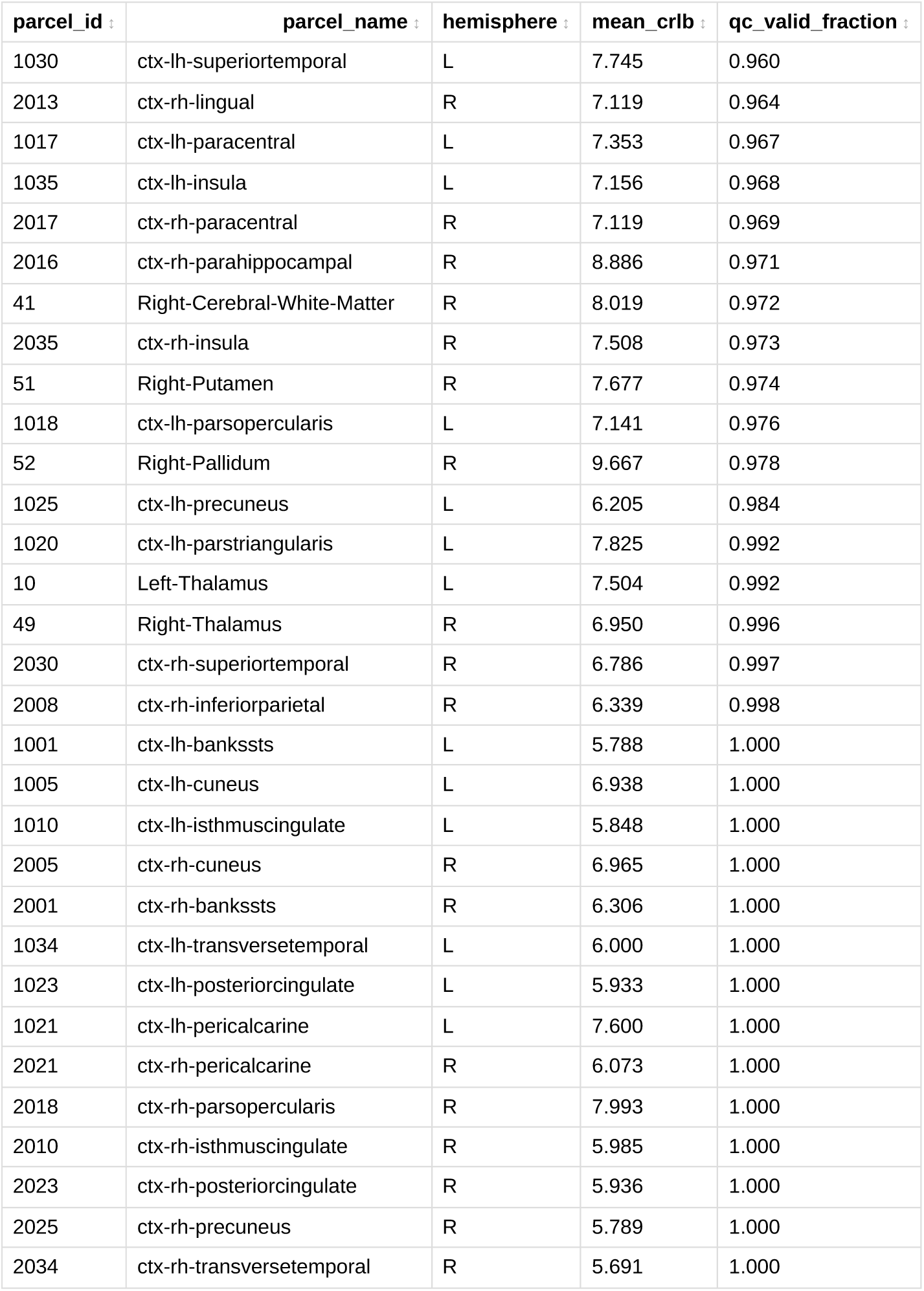

##### Parcellation

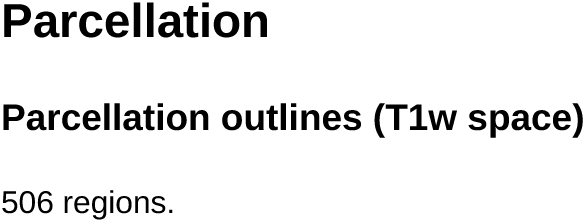
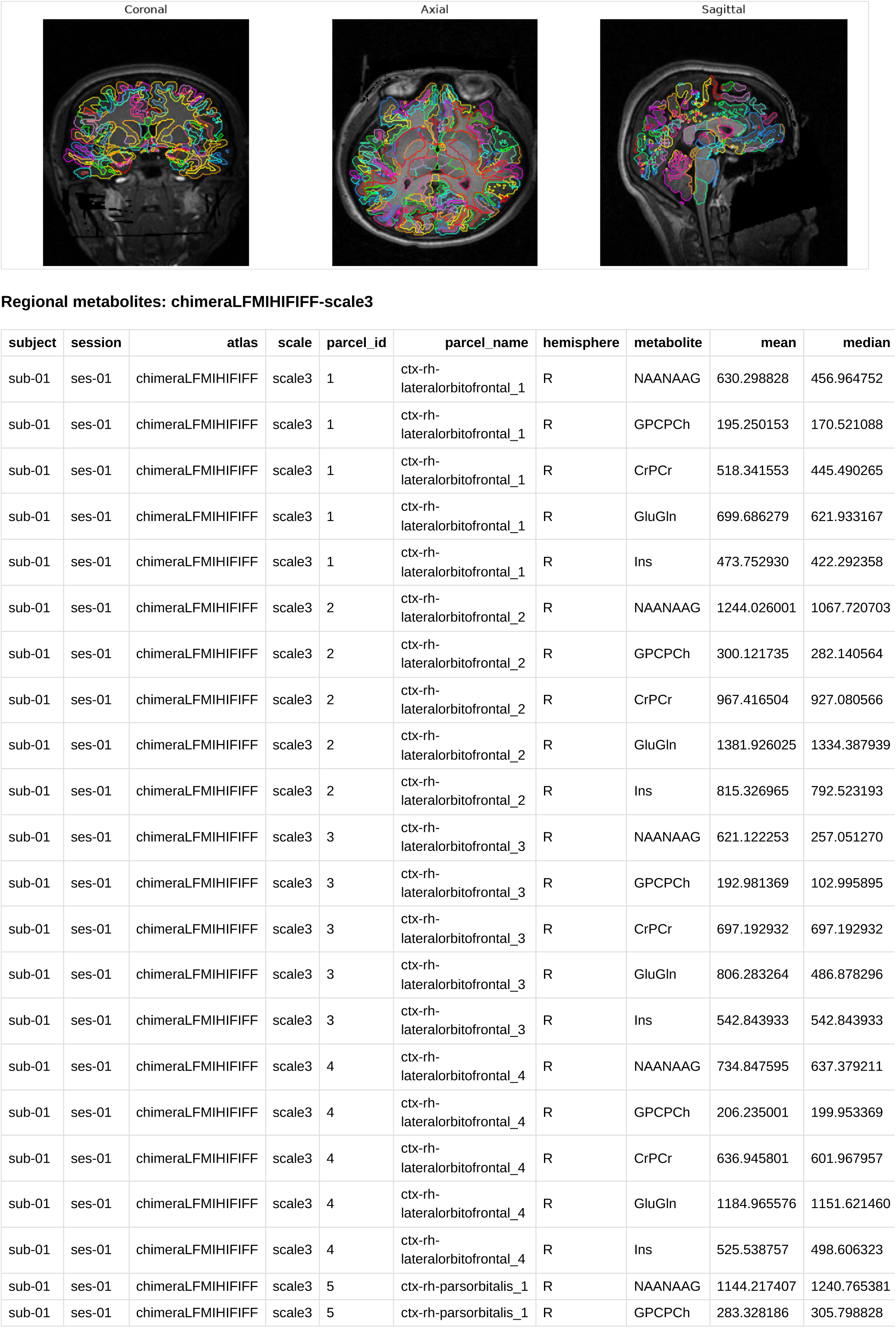
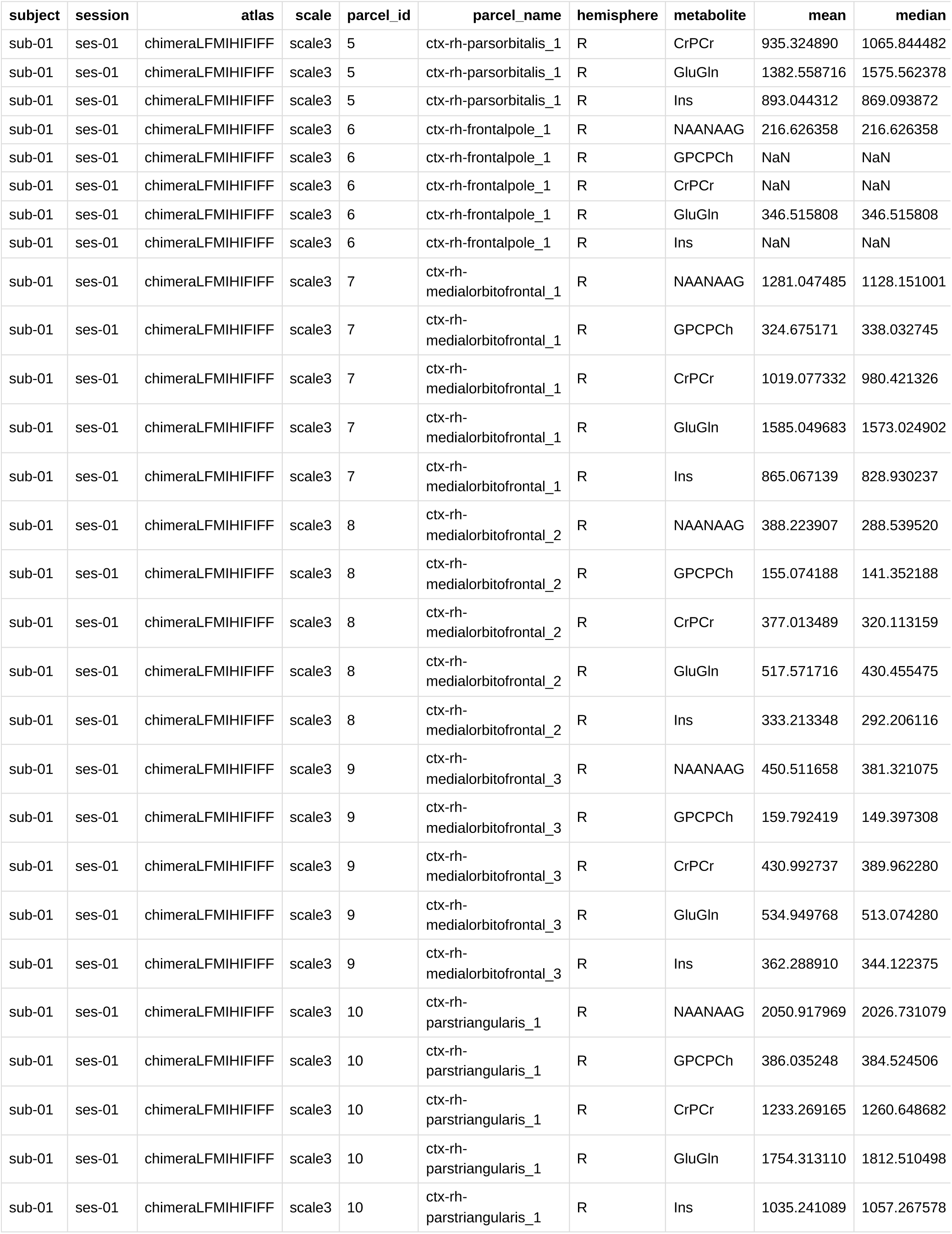

##### Connectivity

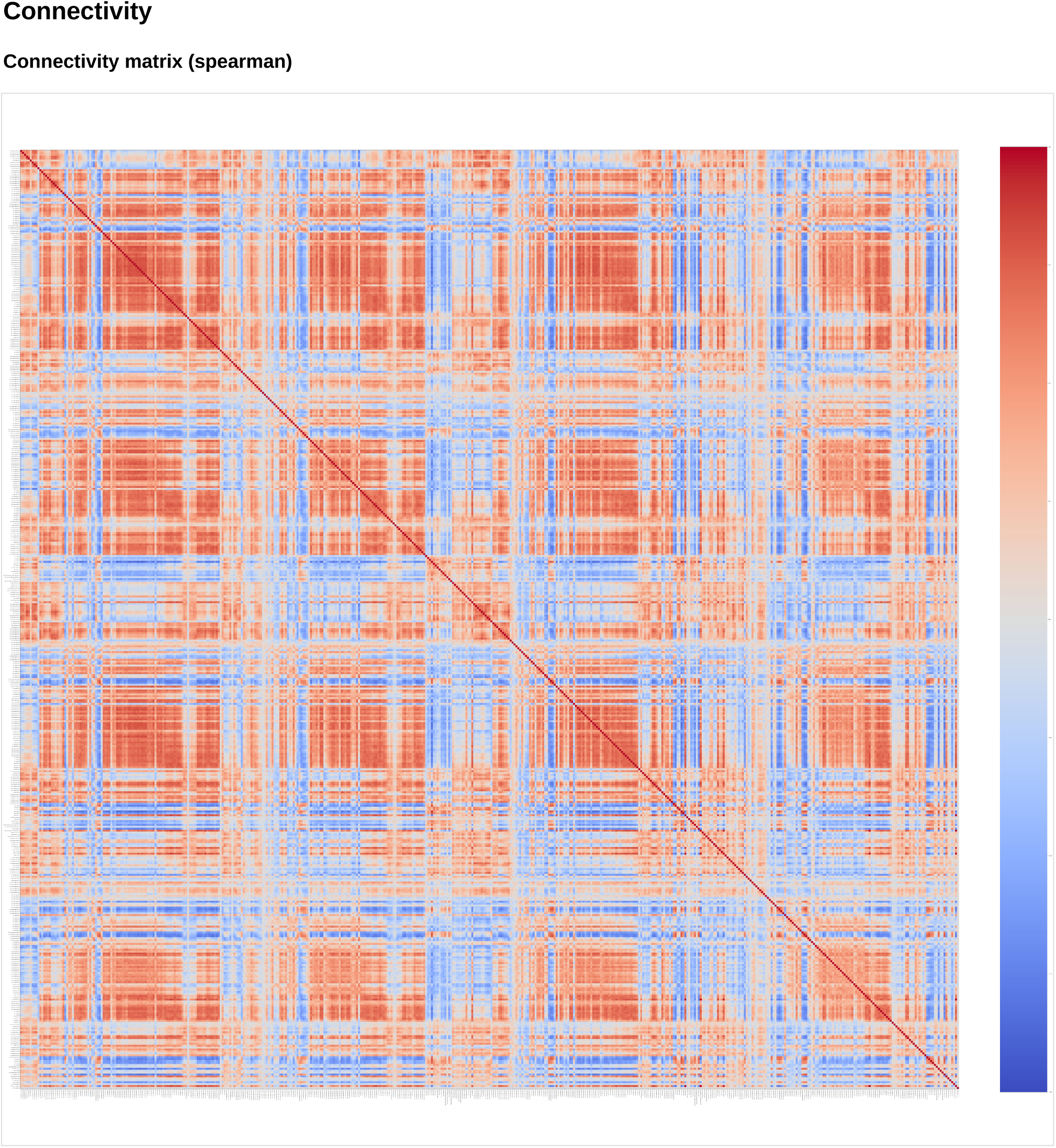

##### MRSinMRS

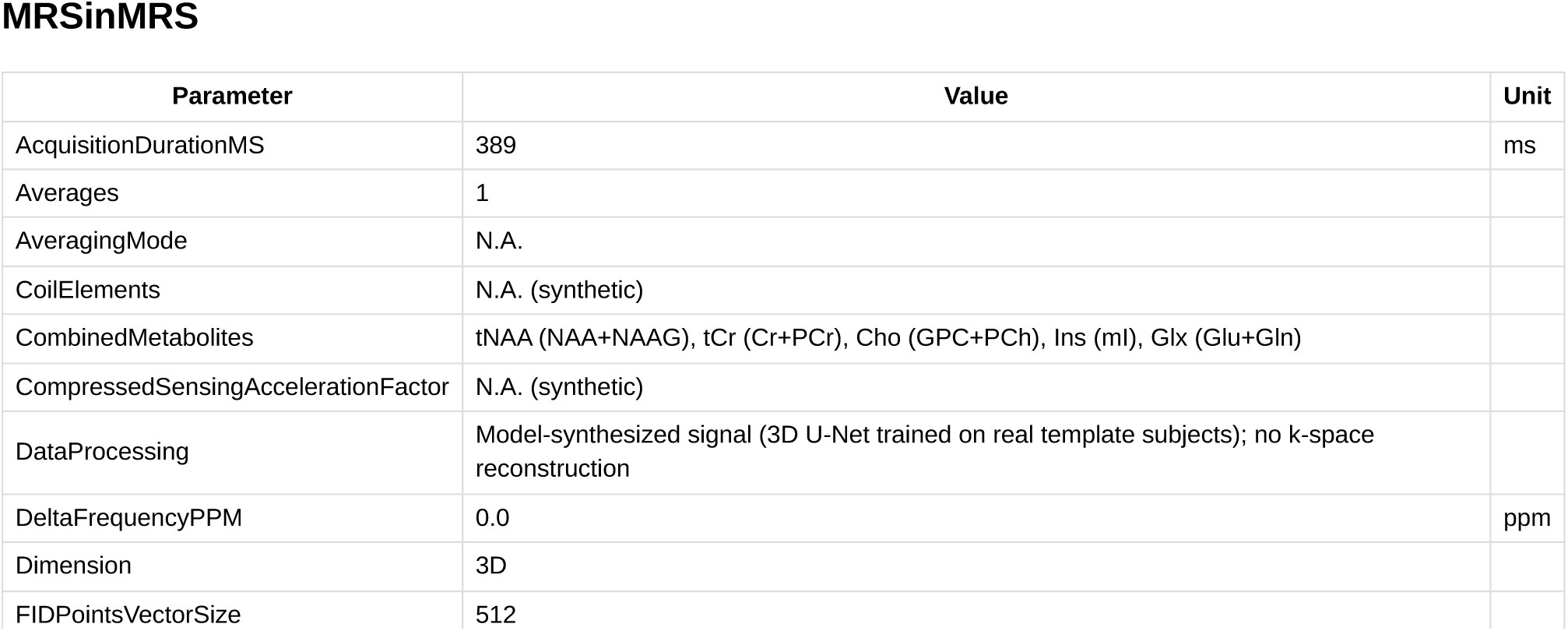
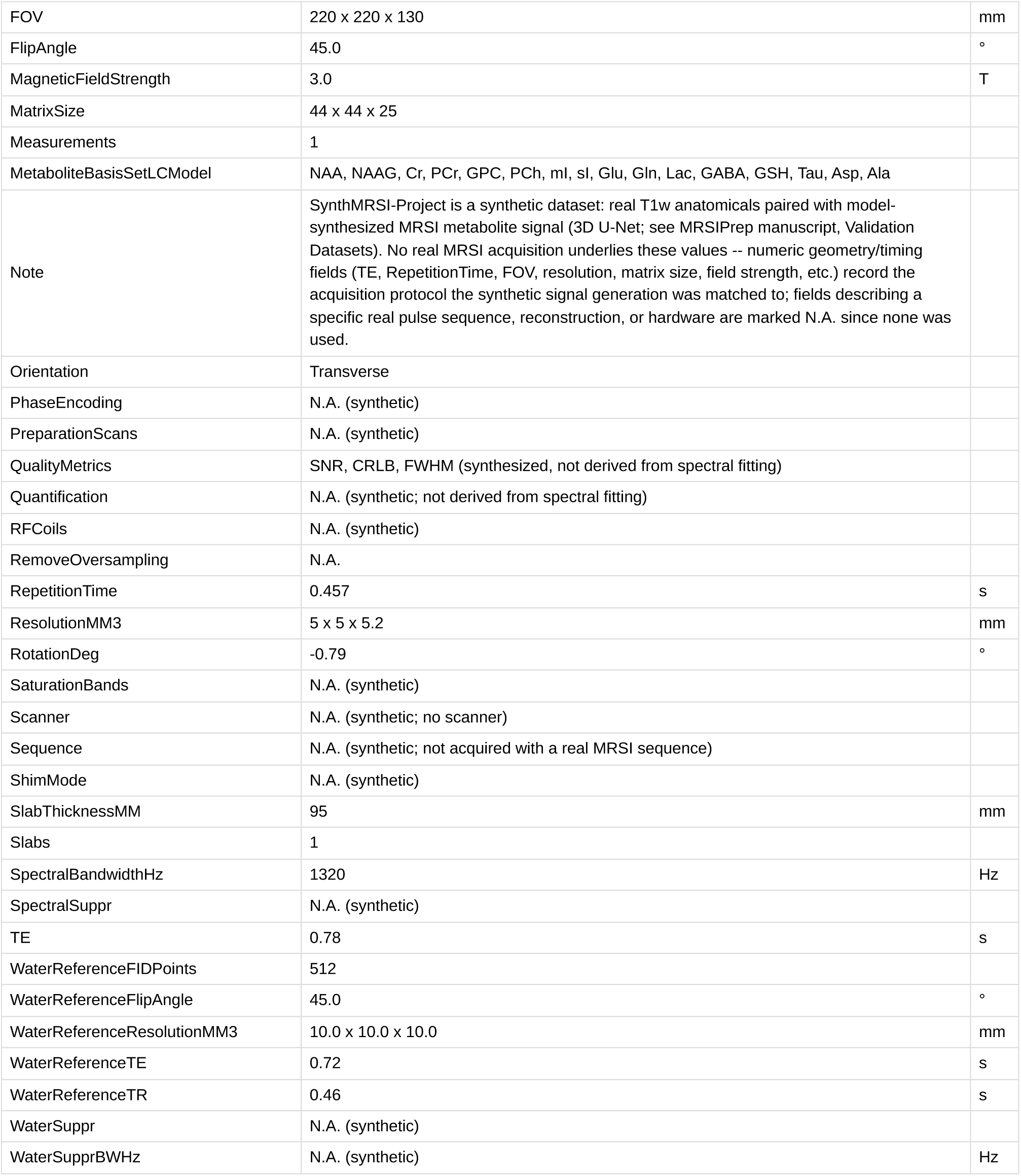

##### PrepParams

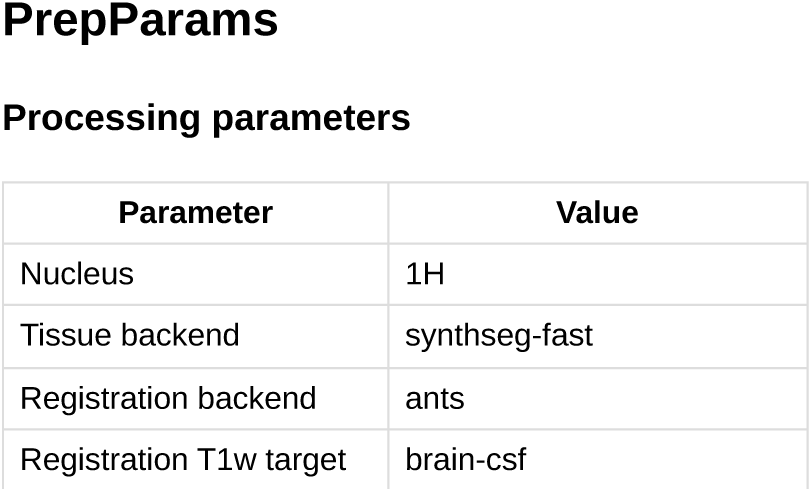
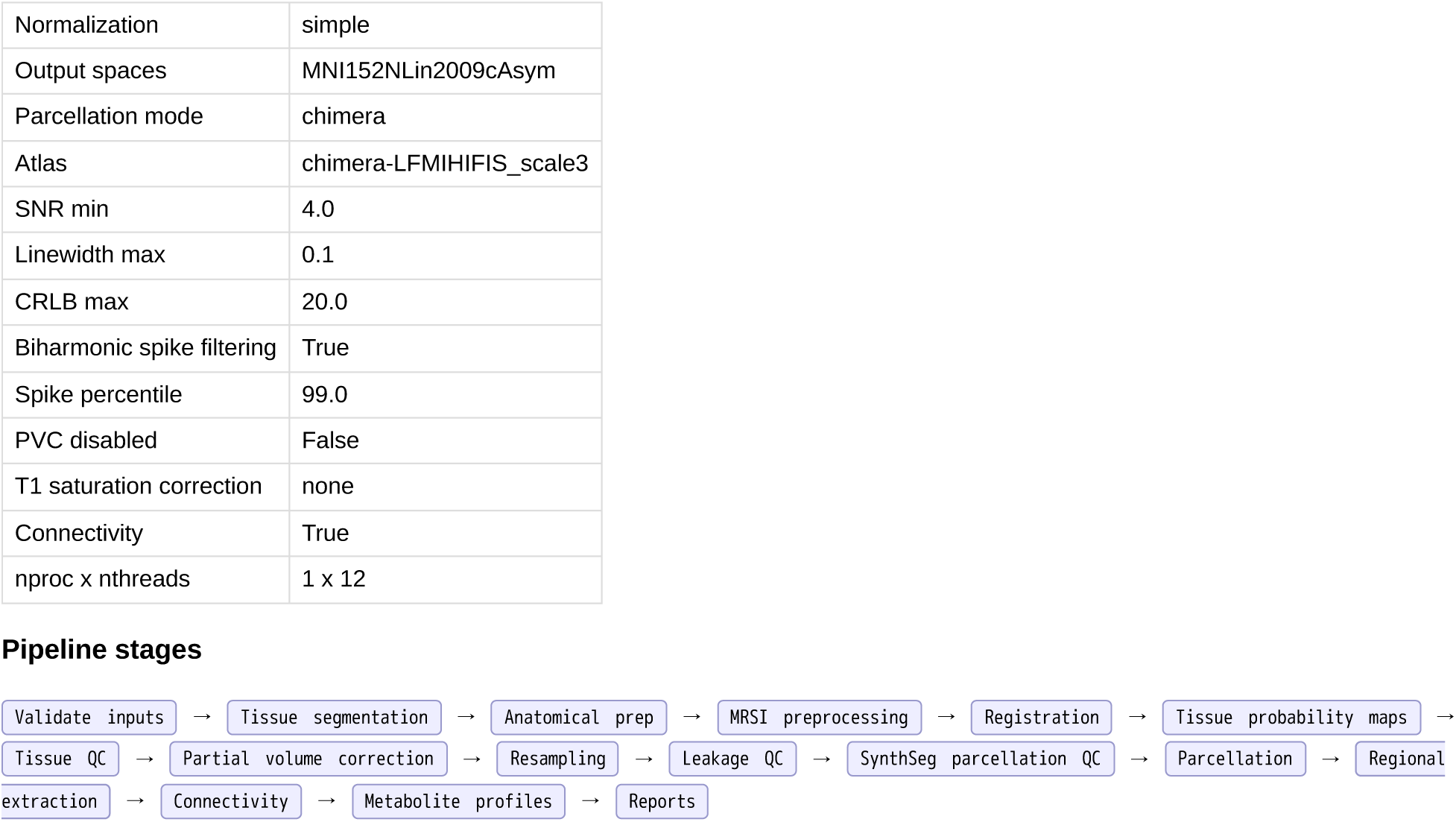

##### Runtime

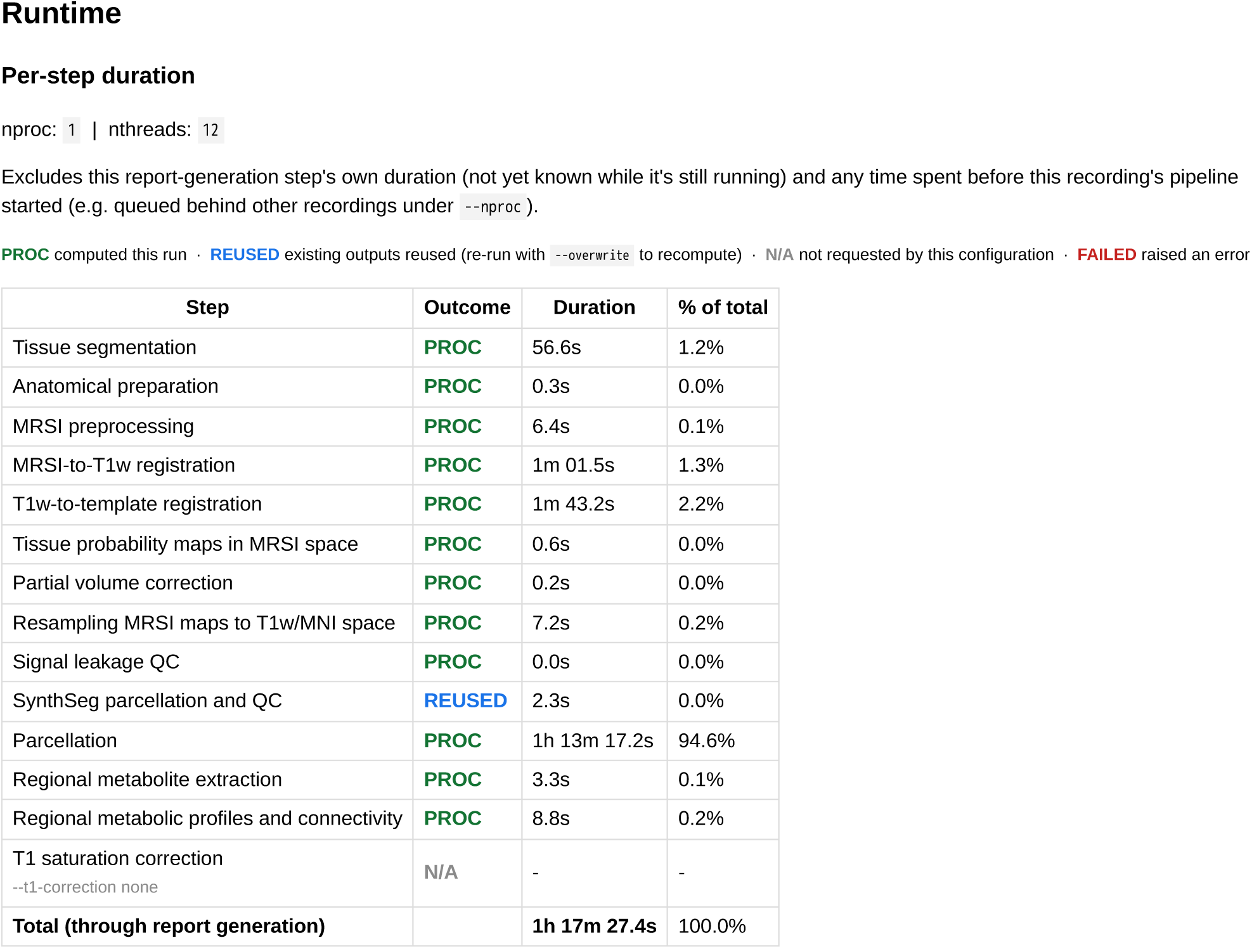

##### Outputs

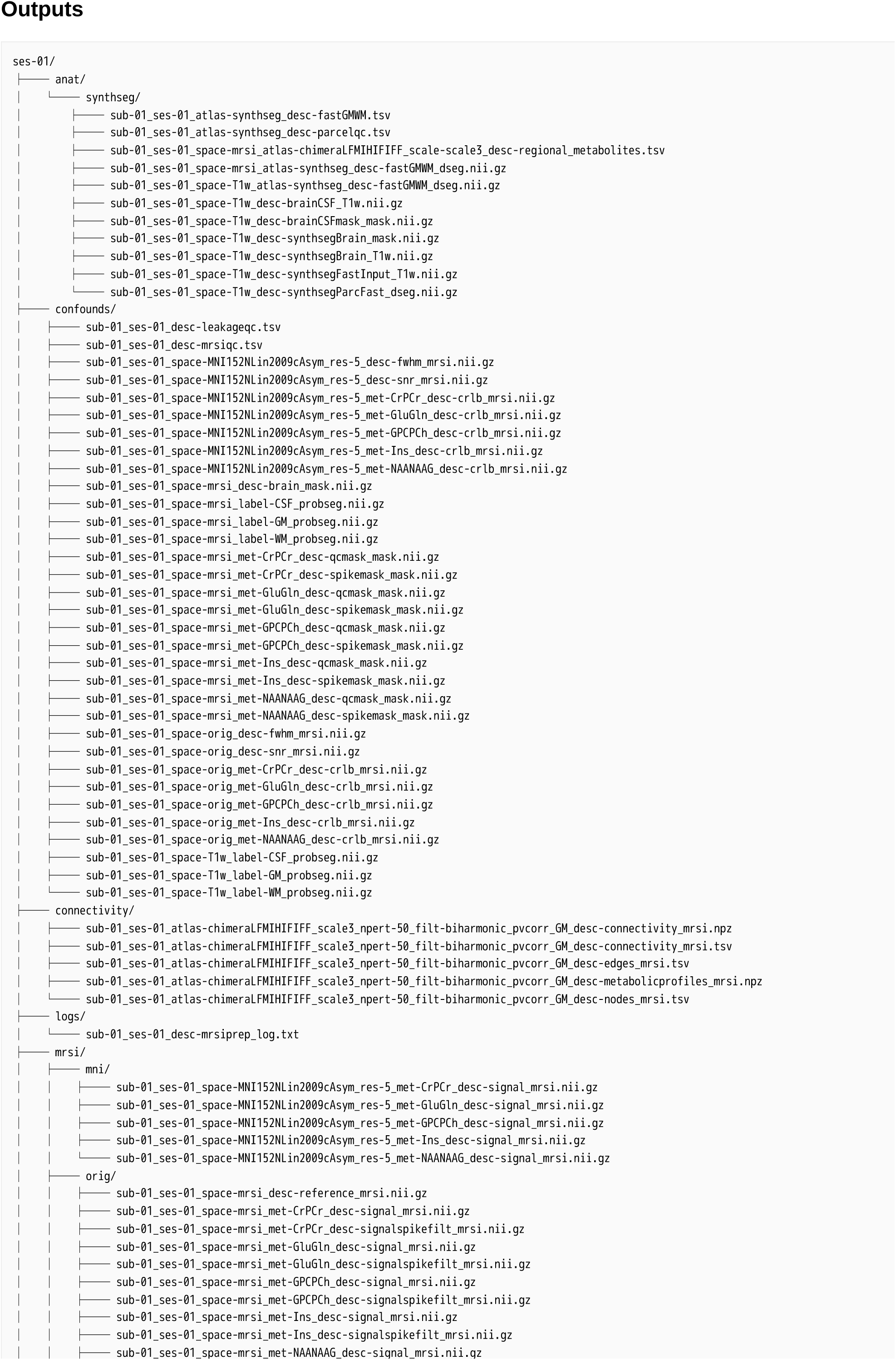
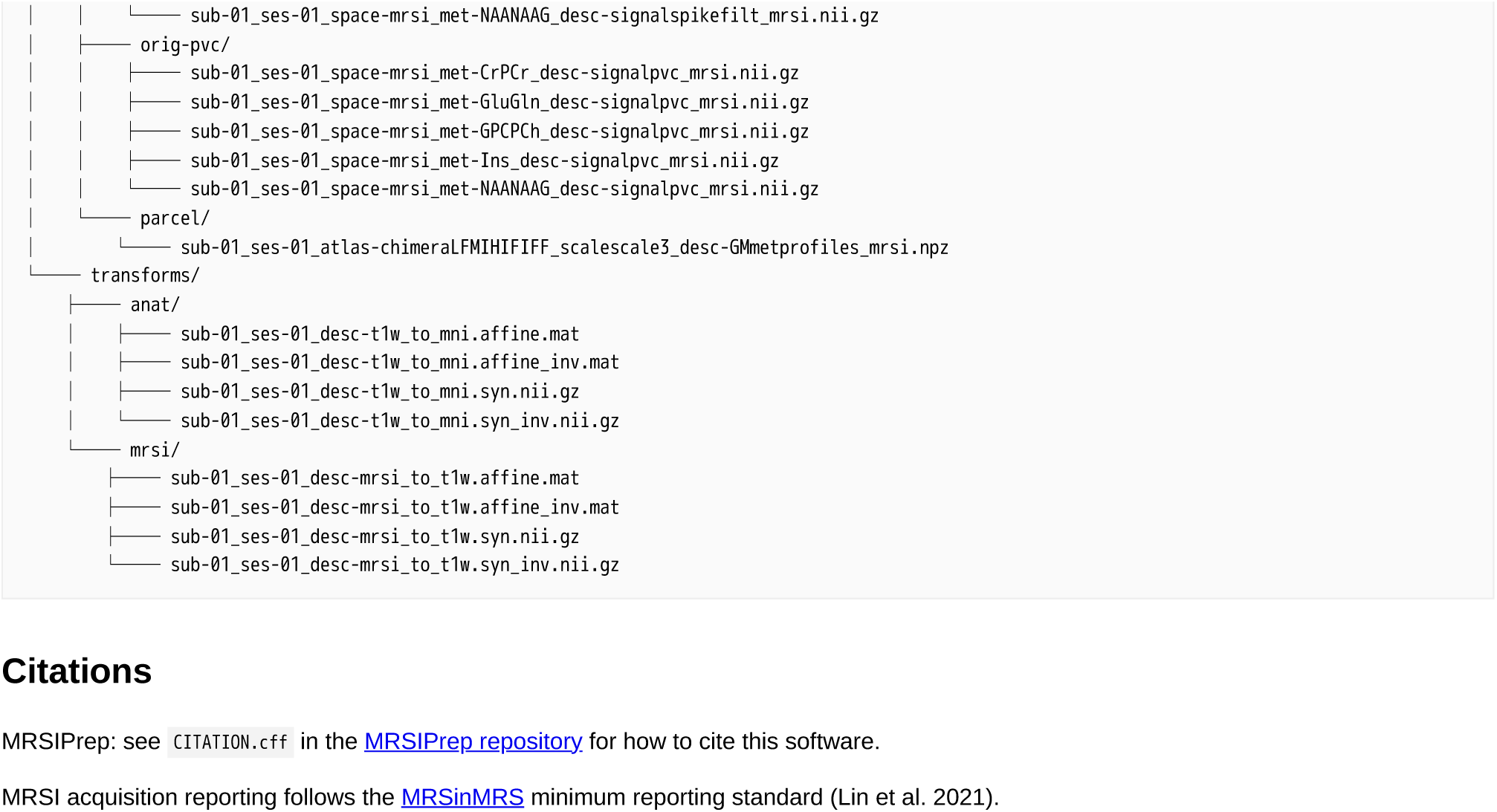

